# Sensitivity of stochastic growth rate and lifetime reproductive output to kin effects: A nonlinear feedback model

**DOI:** 10.64898/2026.10.22.761048

**Authors:** Rahul Mondal, Diego Alburez-Gutierrez, Hal Caswell, Roberto Salguero-Gómez

**Affiliations:** Department of Biostatistics and Epidemiology, International Institute for Population Sciences, Mumbai, India; Max Planck Institute for Demographic Research, Rostock, Germany; Woods Hole Oceanographic Institution, Woods Hole, USA; Department of Biology, University of Oxford, South Parks Road, Oxford, UK

**Keywords:** Kin selection, Lifetime reproductive output, Nonlinear feedback model, Stochastic growth rate, Stochastic sensitivity

## Abstract

Humans exhibit a coevolutionary relationship between the trajectory of social evolution and senescence. A focal individual’s kinship network is a function of his/her life history, while that network in turn shapes the focal individual’s life history through kin effects on reproduction, generating a dynamic, nonlinear feedback loop across the population. Assessing directional selection on kin effects, therefore, requires explicitly modelling the feedback between the kinship network and life history. Here, we develop such a framework and apply it to examine selection pressures on the cooperative behaviour of closely related kin, including mothers, grandmothers, and older sisters. We hypothesise that the negative association of grandmothers, mothers, and older sisters (*>* 10 years) with child mortality are under positive selection because increasing the absolute values of these kin effects would increase the stochastic growth rate. Furthermore, we hypothesise that the positive association of older sisters with child mortality is subject to negative selection, as increasing this effect would reduce the stochastic growth rate. We estimate kin-specific slope coefficients for child mortality using a Bayesian hierarchical model and solve a system of nonlinear equations linking kinship structure and survival to account for the nonlinear feedback between kinship and child survival. Thereafter, we compute stochastic sensitivities of both the stochastic growth rate and the mean lifetime reproductive output to these slope coefficients. Consistent with our hypotheses, all slope coefficients for kin presence on child mortality are negatively sensitive to both fitness metrics. We discuss the applications of our framework to examine the coevolution of social behaviours and senescence in age-structured populations.

**Highlight:**

- A time-varying nonlinear feedback model for the coevolution of life histories and kinship networks is presented.
- The model is valid for any age-structured population.
- The model enables sensitivity analysis of stochastic growth rate and lifetime reproductive output to kin effects on demographic rates.
- An application of the model to multiple countries at different stages of the demographic transition is presented.

## Introduction

The evolution of social behaviours among genetically related individuals can influence the evolution of senescence itself. Kin selection theory predicts that an individual’s inclusive fitness depends on its own reproduction as well as its influence on the reproduction of its relatives through cooperation (Hamilton, 1964). Researchers have argued for decades that the selection for early reproductive cessation and an extension of the post-reproductive lifespan among human females is attributable to the inclusive fitness older mothers gain by providing care to offspring and grand-offspring (Bourke, 2007; Hawkes et al., 1998; Rogers, 1993). Additionally, the pattern of ageing in the focal individual is subject to kin selection, as its longevity can affect the fitness of kin, which in turn, alters the focal individual’s inclusive fitness. In this regard, Lee (2003, 2008) illustrated how the force of selection on age-specific mortality depends on the weighted average of remaining lifetime fertility and the remaining inter-generational transfer of resources at that age. Hence, the interaction between kin selection and senescence is particularly important for humans, as we have evolved to become increasingly dependent on postnatal care for juvenile survival and development of cognitive, linguistic and social abilities (Mace, 2000; Geary and Flinn, 2001; Pavard and Branger, 2012).

Humans exhibit a coevolutionary relationship between the trajectory of social evolution and senescence. Carey and Judge (2001) proposed that selection for parental care and increased investment per offspring (*i*.*e*., intergenerational transfer) leads to greater juvenile survival and reduced parental fecundity, thereby increasing parental survival due to reduced cost of reproduction. This extended parental lifespan provides more opportunities for inter-generational transfer (*i*.*e*., investment per offspring), initiating a self-reinforcing mechanism of lifespan expansion. For instance, increasing overlapping generations as a by-product of increased longevity improves juvenile survival and delays maturity by (*i*) allowing parental support for offspring and (*ii*) reducing the cost of reproduction to mothers through grandmothering (Hawkes et al., 1998). Lucas and Keller (2020) reiterated that lowering extrinsic mortality through extended parental support, coupled with delayed reproduction among young helpers (sisters), slows the pace of life and, in turn, can lead to increasing longevity. However, increasing kin size, especially within the same generation, can also incur costs by increasing competition over resources, thereby offsetting the longevity benefit of kin interaction (Lucas and Keller, 2020). Hence, the reciprocal causality between longevity and the cooperative behaviour of kin could create a positive feedback loop, promoting lifespan extension associated with pronounced kin interaction.

In humans, closely related kin are the primary caregivers who influence the vital rates of the focal individual. In particular, the presence of the mother is associated with increased child survival (Sear and Mace, 2008; Pavard and Branger, 2012). Additionally, empirical evidence suggests that grandmothers have a positive effect on child survival, with maternal grandmothers being more reliable caregivers than paternal grandmothers (Sear and Mace, 2008; Beise and Voland, 2002). The grandmother’s support for the daughter’s offspring is often proposed as a possible solution to the menopause and to selection for post-reproductive lifespan extension, i.e., the grandmother hypothesis (Hawkes et al., 1998). The literature suggests that fathers and grandfathers are substantially less influential than mothers and grandmothers in shaping child survival (Sear and Mace, 2008). Interestingly, mothers may also divide the reproductive labour with their own children, particularly daughters, to reduce the cost of reproduction. Although rare, studies investigating the effect of siblings ‘helpers at the nest’ on child survival show that older sisters have a positive effect on the survival of younger siblings (Sear and Mace, 2008). However, the competition for optimal allocation of the limited family resource complicates the effect of older siblings, sometimes showing that older siblings increase the risk of death for children (Muhuri and Preston, 1991; Lucas and Keller, 2020). Hence, as child survival may be the most important determinant of reproductive success for women (Jones, 2005), examining selection on the caregiving behaviour of closely related kin is particularly important in humans.

Different approaches have examined the coevolution of kin interactions and life-history traits. The main challenge is that the kinship network is a function of life history (Caswell and Song, 2021; Coste et al., 2021; Butterick et al., 2025), which itself is influenced by kin interaction (Pavard and Branger, 2012). Therefore, examining the coevolution of kin interaction and life history requires a dynamic model that accounts for the feedback expressed as a function of life-history traits. Carey and Judge (2001) proposed a theoretical model for a self-reinforcing loop for kin interaction and longevity extension; however, they lacked insight into formal modelling of the process. A major advancement was made by Cyrus Chu and Lee (2006), who developed a formal model using an optimal life history approach to examine the coevolution of lifespan and intergenerational transfer. Researchers have also examined the effects of kin, such as mothers, grandmothers, and siblings, on life history, using multi-state matrix population models in which kin are modelled as additional stage structures (Pavard and Branger, 2012). However, these models do not account for the processes by which the effect of kin interaction cascades through the kinship network, or they treat kin as an extrinsic variable to the model.

Incorporating these diverse kinship effects into the model is challenging because the kinship model, like all structured models, assumes that all individuals experience the same demographic rates (Caswell and John, 1992). Croll and Caswell (2025) approached this by allowing kin effects to apply equally to all individuals, and then solving for the set of demographic rates that both incorporate the effects and produce equal rates for all. Their model was applied to African elephants. Given the data, they perforce had to use a time-invariant model and to rely on ad hoc methods to incorporate reports of the effects of kin on juvenile survival and female fertility.

In our analysis, we follow the Croll and Caswell approach, but develop it in a time-varying context and implement a Bayesian procedure to provide rigorous estimates of kin effects directly from demographic data. Our model makes it possible to examine the sensitivity of two measures of fitness to the perturbation in kin effects: stochastic growth rate and mean lifetime reproductive output (LRO, hereafter), for population-level and cohort-level inference, respectively. We use this framework to test two specific sets of hypotheses. First, the negative association of the presence of (H1a) mother, (H1b) grandmother, and (H1c) older sister aged *>* 10 years with child mortality would be under positive selective pressures. Increasing the absolute magnitude of the (negative) kin dependence would increase the stochastic growth rate and the mean lifetime reproductive output. Our expectation is based on studies showing how the stochastic growth rate is positively affected by the presence of the mother and grandmother in child survival (Metcalf and Pavard, 2007; Pavard and Branger, 2012). Second, (H2) the positive association between the presence of an older sister aged ≤ 10 years and child survival will be subject to negative selective pressures. Older siblings acting as competitors for resources may dilute the positive effects of intergenerational transfer of resources and care, thereby under negative selection pressure (Lee, 2003; Lucas and Keller, 2020). Our framework contributes to the study of kin selection and the evolution of senescence by integrating dynamic feedback in kin relationships to perform perturbation analyses of the stochastic growth rate and mean lifetime reproductive output, accounting for the nonlinear feedback between the kinship network and life-history traits.

## Methods

To examine the sensitivity of stochastic population growth and lifetime reproductive output (LRO) to the kin dependence, we developed a quantitative framework comprising seven steps. (1) We construct matrix population models based on survival and fertility matrices for the examined countries and years (panel A in Figure 1). (2) Using the survival and fertility matrices, we construct kinship networks by the age of the focal (panel B in Figure 1). (3) We then parameterise the survival matrix as a function of the kinship network using a Bayesian hierarchical model (panel C in Figure 1). (4) We account for the nonlinear feedback between kinship network and survival matrices by solving a set of nonlinear equations (panel D in Figure 1). (5) Using the solved matrix models, we computed the long-term stochastic growth rate and mean LRO (panel E in Figure 1). (6) To test our hypotheses H1 and H2, we compute the sensitivity of the stochastic growth rate and mean LRO to kin dependence (panel F in Figure 1). All analyses were performed in R version 4.5.3 (Team, 2025).

**Figure 1:**
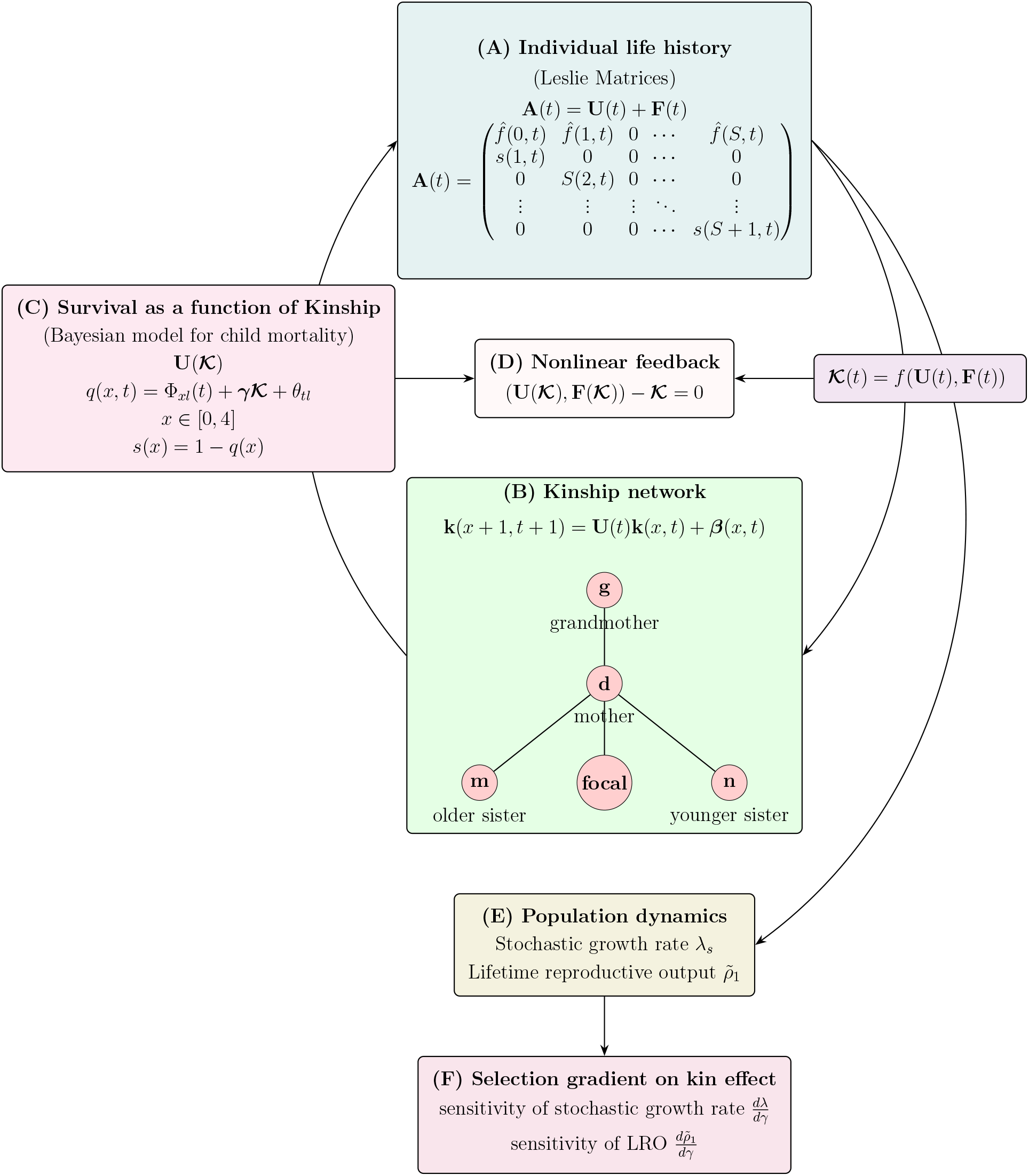
Schematic diagram of the methodological framework. The figure illustrates the quantitative pipeline for examining the selection gradient on kin effect across six major steps. (A) First, we construct the Leslie matrices from survival and fertility rates. (B) Using the survival and fertility rates (matrices), we construct the kinship network. (C) We then model child survival as a function of the kinship network using a Bayesian hierarchical model. (D) Next, we solve the nonlinear feedback between kinship and survival matrices to find the kinship network and survival rates that exactly reproduce each other. (E) Then, using the solved survival matrices, we construct the Leslie matrices and compute the stochastic growth rate and mean lifetime reproductive output. (F) Finally, we examine the sensitivity of stochastic growth rate and mean lifetime reproductive output to the slope coefficient of the kinship network on (child) survival.

### Matrix Population Models

To test our hypotheses (H1 and H2) regarding stochastic sensitivity of fitness metrics to kin dependence, we first constructed matrix population models for six countries. Specifically, we considered a population that grows according to the following equation:

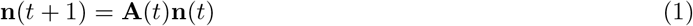

where **A**(*t*) is the projection matrix at time *t* (Caswell, 2001). For an age-structured human population, where individuals range between ages *x* = 1, …, *ω*, the matrix **A**(*t*) is the Leslie matrix at time *t* comprised of the survival (***U*** *(t)*) and fertility (***F*** *(t)*) matrix as shown below:

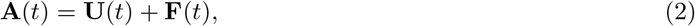

or

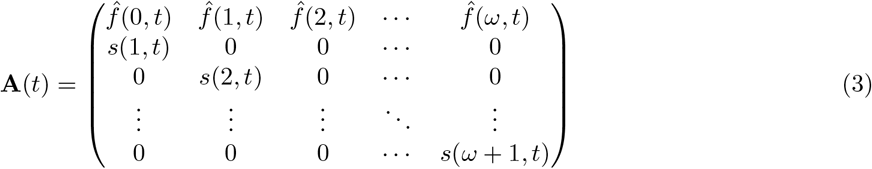

(Leslie, 1945, 1948). The survival matrix **U**(*t*) is an *ω × ω* matrix where subdiagonal entries represent the probability of surviving from age *x* to *x* + 1 during time *t* to *t* + 1 and all other entries are zero. The *ω × ω* matrix **F**(*t*) has age-specific reproductive rates in the first row obtained as

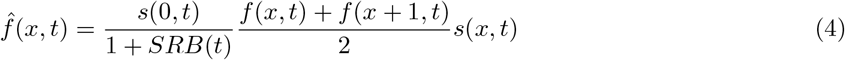

where, *s*(0, *t*) is the infant survival rate, *s*(*x, t*) are age-specific survival rates given in the subdiagonal elements of **U**(*t*), *f* (*x, t*) is the age-specific fertility rates and *SRB*(*t*) is the sex-ratio at birth at time *t* (Preston et al., 2001). Following the standard demographic convention for constructing one-sex MPMs, we set *SRB* to 1.05 for all countries and years (Wheldon et al., 2013). All other elements of the fertility matrix are zeroes. Here, we constructed the one-sex (*i*.*e*., female-dominant) model for **U** and **F** using complete period life tables.

To examine how the selection gradient on kin dependence varies across socio-demographic environments, we strategically selected six countries: two countries each from three distinct stages of demographic transition. The Sub-Saharan countries of Ethiopia and Gambia represent “*type-1*” populations at an early stage of demographic transition, with a total fertility rate (TFR, hereafter) of ~4, a life expectancy of ~67 years, and a mortality profile that is predominantly characterised by communicable, maternal, neonatal, and nutritional (CMNN, hereafter) disorders (United Nations, 2024; Vollset et al., 2024). Although the Ethiopian and Gambian populations cannot be characterised by natural fertility and natural mortality as in hunter-gatherers, child mortality is considerably high. Second, the South Asian countries of Bangladesh and India represent “*type-2* “ populations, situated in the context of declining fertility (TFR ~ 2) and improving survival (life expectancy ~ years), characterised by a shrinking but noticeable share of CMNN and increasing share of Non-Communicable diseases [NCDs, hereafter; (United Nations, 2024; Vollset et al., 2024)]. Finally, Japan and Sweden represent the “*type-3* “ population at the advanced stage of demographic transition, with a very low TFR (~ 1) and under-five mortality of ~ 0.002, such that the role of extrinsic factors in child mortality is very low (United Nations, 2024; Vollset et al., 2024). As “*type-2* “ countries are transitioning to the advanced stage of demographic transition and “*type-3* “ countries are at the advanced stage, modelling the baseline mortality that is unaffected by kinship becomes crucial to minimise false positives (Sear and Mace, 2008).

To construct the MPMs for selected countries, we collected data on complete period life tables and fertility rates from the World Population Prospects 2024 (United Nations, 2024). Specifically, we extracted the age-specific survival rates *s*(*x, t, l*) from the complete period life table for females in ages *x* = 0, 1, …, 110+, years *t* = 1950, …, 2023 and countries *l* = 1, …, 6, and the age-specific fertility rates *f* (*x, t, l*) for ages *x* = 13, 14, …, 55+ and examined country-year. Using the survival and fertility rates, we constructed the MPMs through (3) and (4).

### Kinship Model

To examine the effect of kin interactions on stochastic growth rate, we constructed kinship networks. Specifically, we used the time-varying kinship model (Caswell and Song, 2021) to compute the expected number of kin of type **k**(*x, t*) at age *x* of the focal individual at time *t*, considering time-varying vital rates whose dynamics are given by

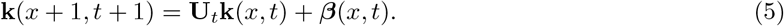

Here, the subsidy vector ***β***(*x, t*) is the age structure vector of the new kin arriving between *x* and *x* + 1. The vector ***β***(*x, t*) either takes the form

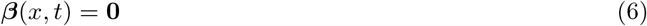

if there are no new kin of type **k**, or

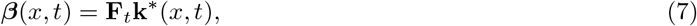

which applies the fertility **F**(*t*) at time *t* to the age structure vector of another kin **k**^∗^ that produces offspring of type **k**. Moreover, we defined a set of boundary conditions, *i*.*e*., the complete age vector **k**(*x*, 0)at time *t* = 0, and the initial age vector **k**(0, *t*) at each time *t* (as in Caswell, 2019a; Caswell and Song, 2021). We computed **k**(*x*, 0) using the **U** and **F** for the year 1950, with the assumption that **U**_1950_ and **F**_1950_ has been operating for a long time. To calculate **k**(0, *t*), we considered

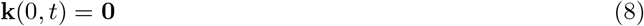

if the focal individual has no kin of type **k** at her birth, and

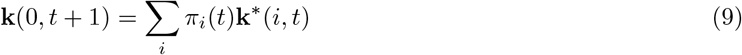

if the focal individual has possible kin of type **k** at birth. Here, *π*(*i, t*) is the proportion of mothers who reproduce at age *i* at time *t* obtained as

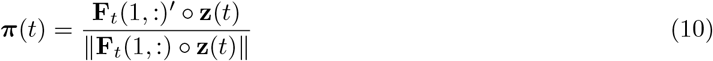

where **z** is the age-structure vector (see Caswell, 2019a; Caswell and Song, 2021). Thus, the kinship model (5) produces the expected number of kin in the kin network

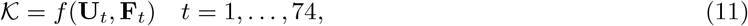

depending on survival **U**_*t*_ and fertility matrix **F**_*t*_ for the 74 years (*t*) between 1950-2023.

To examine the effects of grandmothers, mothers, and older sisters on the stochastic growth rate and LRO, we computed expectations of these kin. Note that when we refer to the kinship network, we refer to this subset of the total network. Hence, we computed the expected number of mothers as

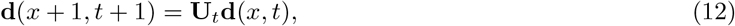

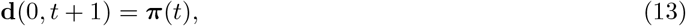

grandmothers as

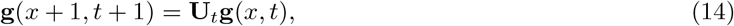

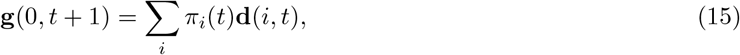

and older sisters as

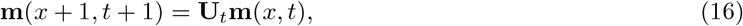

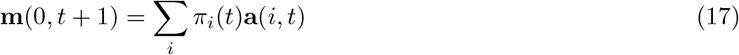

(Caswell and Song, 2021). It is worth noting that the subsidy term ***β***(*x, t*) is zero and therefore not present in (12), (14), (16) because at birth of focal they can not gain a new grandmother, mother, and older sisters. Therefore, the projected set of kinship networks is

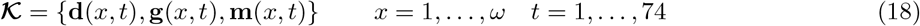

We implemented the models to obtain the expected number of mothers **d**(*x, t*), grandmothers **g**(*x, t*), and older sisters **m**(*x, t*) using the DemoKin package version 1.0.3 in R (Williams and Alburez-Gutierrez, 2025).

The set *K* is large. Each of the three entries is itself a 3-dimensional array. For example, **d** contains the abundance of *ω* age classes of mothers at each of *ω* age classes of Focal at each of *t* = 1, …, 74 time points, with the same structure for **g** and **m**. This amounts to 895,400 entries for each type of kin.

### Feedback Model

To model the feedback of the kin network *K* into the survival rate, we expressed the survival matrix as a function of the kin network **U**(*K*). We developed the feedback model in four steps. As in Croll and Caswell (2025), we expressed the kinship network in a set of nonlinear equations

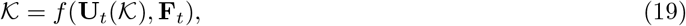

or

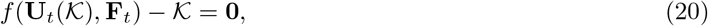

where the survival matrix **U**_*t*_(*K*) is a function of the kinship network *K*, which itself is a function of the time-series of the survival matrices **U**_74_, …, **U**_1_.

As a link between the kinship network and the survival matrix, we use the child mortality probability *q*(*x, t, l*) for age *x* ∈ [0, 4] and country *l* = 1, …, 6. We model this mortality as a function of kin using a Bayesian hierarchical model, explained in the subsequent section. Thereafter, we obtained the kinship network 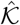 by numerically solving (20) using the nleqslv package version 3.3.7 in R (Hasselman and Carnell, 2026), where the difference between the computed kinship network *K* and the solved kinship network 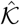 is minimised using a Newton gradient search method. Finally, we computed the corresponding survival matrices 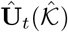, the fertility matrices 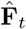, and the Leslie matrices at time *t* as

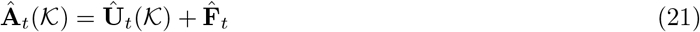

representing a kin-dependent population model.

### Bayesian Model for Kin Interaction

To parameterise the child survival as a function of the kinship network, we developed a Bayesian hierarchical model. Specifically, we model the probability of death *q*_*xtl*_ for age *x* ∈ [0, 4], year *t*, and country *l* as

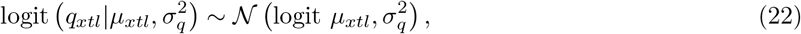

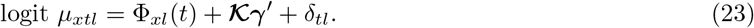

We formulated the model in (23) to estimate the global slope vector ***γ*** of the three kin in *K* for child mortality, while accounting for baseline mortality. Here, *µ*_*xtl*_ is the latent child mortality, the term Φ_*xl*_(*t*) represents the age-country-specific smooth temporal trend in mortality, and *δ*_*tl*_ is the random country-specific temporal fluctuations in mortality, together accounting for the baseline mortality unaffected by the kin *K*. Note that we assumed the same slope coefficient of kinship on child mortality across all six countries, after accounting for country-specific baseline mortality and random fluctuations in mortality.

To model the baseline smooth temporal trend in mortality, we defined the parameter Φ_*xl*_(*t*) as a linear combination of *P* (*p* = 1, …, *P*) B-spline basis functions **B**_*p*_(*t*)

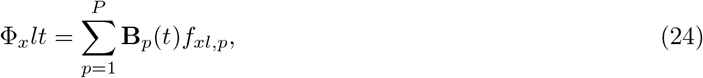

where *f*_*xl,p*_ is the spline base coefficient of knot *p*, age *x*, and country *l*. We used a cubic B-spline with equally spaced knots with a between-knots interval length of 2.5 years (as in Alkema and New, 2014). Thereafter, we penalised the second-order difference in adjacent spline coefficients (Δ^2^*f*_*xl,p*_ = *f*_*xl,p*_ − 2*f*_*xl,p*−1_ + *f*_*xl,p*−2_)?.

We do so by modelling the spline coefficient as

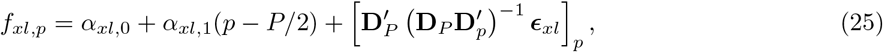

where *α*_*xl*,0_ and *α*_*xl*,1_ are the level and slope parameters, respectively, for the spline coefficients in age *x*, and country *c*. The parameter ***ϵ***_*xl*_ = (*ϵ*_*xl*,1_, …, *ϵ*_*xl,P* −2_) contains *P* − 2 the second-order difference in spline coefficients, and **D**_*K*_ is the difference matrix defined as

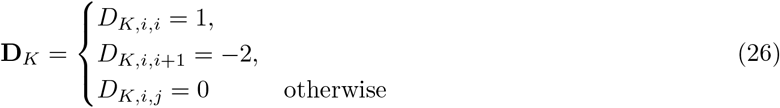

(Currie and Durban, 2002; Eilers and Marx, 2010; Alkema and New, 2014). Finally, we penalised the second-order differences in spline coefficients by imposing

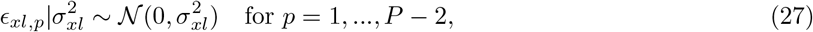

where the variance 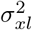 determines the extent of smoothing (as in Alkema and New, 2014). Note that penalising the second-order difference between the adjacent spline coefficients *f*_*xl,p*_ and computing the baseline mortality as (24) gives us a smooth temporal trend of mortality for each age *x*, and country *l*.

To model the association of kinship on child mortality, we included kin availability as a covariate. Specifically, we used the probability of having a living mother *d*(*x, t, l*), grandmother *g*(*x, t, l*), older sister aged 10 year or less *m*_≤10_(*x, t, l*), and older sister aged above 10 years *m*_*>*10_(*x, t, l*) for focal aged *x* in year *t* and country *l, i*.*e*., ***K*** = *{d, g, m*_≤10_, *m*_*>*10_*}*. Because individuals can have exactly one mother and one (maternal) grandmother, the probability of having a living mother and a living grandmother follows a binomial distribution. Thus, we computed the probabilities from the expected number of mothers and grandmothers obtained from the kinship model as

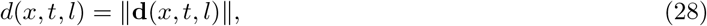

and

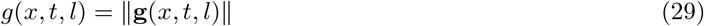

(Croll and Caswell, 2025). Thereafter, considering the probability distribution of the number of older sisters having a Poisson distribution, we computed the probability of having older sisters aged ≤ 10 and *>* 10 as

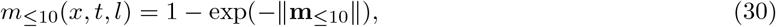

and

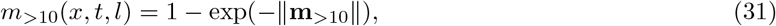

respectively, where **m**_≤10_ and **m**_*>*10_ are the expected number of older sisters obtained from the kinship model (Caswell, 2024; Croll and Caswell, 2025). Finally, we defined the global slope coefficient vector ***γ*** on each covariate in ***K***.

To account for the sharp rise in child mortality due to conflicts or epidemics, we incorporated a country-year-specific random parameter *δ*_*tl*_. Among the examined countries, Bangladesh shows a sharp spike in mortality in 1971, *i*.*e*., the excess mortality attributable to the Bangladesh Liberation War (Curlin et al., 1976), and a smooth declining trend in the other years. Similarly, in Ethiopia, we observed a pronounced peak in the mortality curve between 1960 and 1990 (United Nations, 2024), followed by a smooth curve thereafter. The smooth temporal trend in mortality modelled by the penalised spline term Φ_*xl*_(*t*) can not produce these sharp spikes in mortality. Therefore, we incorporated a country-year-specific random parameter *δ*_*tl*_ in the model (23) to account for within-country temporal fluctuations, defined with a shrinkage prior detailed in the subsequent section.

To perform Bayesian estimation of child mortality in (23), we defined informative priors for most model parameters. Specifically, we defined informative priors on the slope coefficients *γ* by pooling the estimates of slope and standard deviation from studies by Jamison et al. (2002) and Sear and Mace (2008, 2022) using an inverse-variance weighting scheme. The informative prior on the slope coefficient of the kin presence on child mortality improves identifiability of the slope parameters *γ*. In addition to the smooth temporal trend in baseline mortality Φ_*xl*_(*t*), we allowed for sharp spikes in the child mortality by defining a regularised horseshoe prior on the country-year-specific random parameter *δ*_*tl*_. The regularised horseshoe prior shrinks the parameter toward zero, maintaining the time-smoothed baseline mortality, and allows the coefficient to deviate from zero only when the data are sufficiently informative to suggest a spike in the mortality curve. Detailed formulations of priors for each parameter are given in the appendix. Finally, we perform the inference on the model parameters using No-U-Turn Sampler (NUTS), a variant of Hamilton Monte Carlo sampling algorithm (Hoffman and Gelman, 2014) implemented via cmdstanr(Gabry et al., 2025). We provided the detailed specifications for parameter estimation and model diagnostics in the appendix.

### Stochastic Population Growth

Having estimated a temporal sequence of survival and fertility matrices that incorporate the effects of selected kin on child mortality, we can extend the model to compute a stochastic growth rate to test our hypotheses (H1 and H2) regarding the sensitivity of population growth to kin interactions. Specifically, we computed the stochastic population growth rate log *λ*_*s*_ considering the projection model (2) where the Leslie matrix **A**(*t*) is generated by an independent and identically distributed (i.i.d., hereafter) environment that produces, for every environmental state, a set of vital rates that satisfy certain regularity conditions.

We have performed the stochastic growth rate estimation in five steps. First, assuming that each year-specific matrix **A**_*t*_ is a realisation of the environmental state at year *t*, and that the environmental state is generated by an i.i.d. environmental process, we simulated a time series of Leslie matrices for *T* = 1000 years. Here, the Leslie matrix **A**(*r*) at time step *r* = 1, …, *T* is chosen with

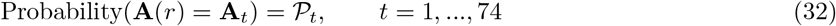

such that **A**(*r*) = **A**_*t*_, if at time-step *r* the environment is in a state represented by the conditions of year *t*. In other words, we sampled a sequence of *r* = 1, …, 1000 matrices **A**(*r*) from the set of matrices *t* = 1, …, 74 matrices **A**_*t*_ using a random sampling with replacement. Hence, in our computation, the probability *P*_*t*_ is considered as a constant *P* = 1*/*74 for all *t*. Second, using the simulated sequence of Leslie matrices **A**(*r*) = (**U**(*r*) + **F**(*r*)) over *r* = 1, …, *T* time-steps we produced corresponding kinship network *K*(*r*) and then computed the probability of having a surviving mother, grandmother, and older sister using equations (28)-(31).

Third, to make the child survival rate kin-dependent, we reexpressed the stochastic sequence of Leslie matrices as **A**^∗^[**Θ**, *K*(*r*)] = (**U**^∗^[**Θ**, *K*(*r*)] + **F**^∗^(*r*)) by substituting

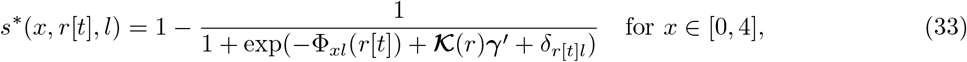

in (3) and (4), where **Θ** = {Φ, ***γ***, *δ}* is the posterior of parameters extracted from (23). It is important to note that we use the year-specific parameters in **Θ** for year *t* to reconstruct survival rates for time-step *r* if **A**_*t*_ is sampled for time step *r, i*.*e*., **A**_*t*_ = **A**(*r*). We denote the use of the parameters for year *t* to estimate survival rates at time step *r* by *r*[*t*]. Here, we constructed the median of **A**^∗^(*r*) using the median of the posteriors of **Θ** in (33). The posterior draws of the parameters from (23) can be used to compute posteriors of **A**^∗^(*r*), indicating the uncertainty in the parameters translating into uncertainty around **A**^∗^(*r*). However, we present the result for the median of the posteriors as the estimation using each posterior draw is computationally very expensive.

Fourth, we solved the nonlinear equation (20) to obtain the Leslie matrices 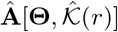 over the simulated time-steps *r* = 1, …, *T*. Fifth, we computed the stochastic population growth rate as

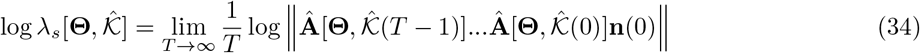

(Tuljapurkar, 1990). We computed the stochastic growth rate using a numerical simulation, as a closed-form analytical solution for (34) is not available.

### Growth Rate Sensitivity

To test our hypotheses (H1 and H2) regarding the selection gradient on kin dependence, we computed the stochastic sensitivity. Specifically, we examined the sensitivity of log *λ*_*s*_ to the parameters **Θ** that determine the Leslie matrices **Â** (*r*) by computing the derivatives of log *λ*_*s*_ with respect to the parameters *θ*_*r*_ at each time step *r*. We computed the derivatives as

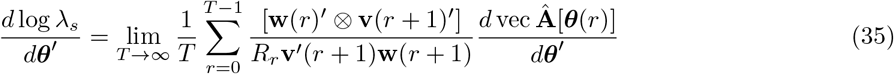

where the vectors **w**(*r*) and **v**(*r*) are the stochastic analogue of the right and left eigenvector corresponding to the dominant eigenvalue of the **Â** (*r*), and *R*(*r*) is the growth rate of total population size from *r* to *r* + 1 (Tuljapurkar, 1990; Caswell, 2019b). If the parameter ***θ*** consists of vital rates elements from **Â**, then *d* vec **Â** */d****θ***^*′*^ = **I**, where **I** is the identity matrix. On the other hand, when ***θ*** consists lower level parameters, *i*.*e*., the slope coefficient vector ***γ*** of kin elements in (23) then the *d* vec **Â** */d****θ***^*′*^ contains the derivatives of **Â** with respect to these parameters (Caswell, 2019b).

### Lifetime Reproductive Output

To examine the sensitivity of mean LRO to kin presence, we computed the LRO assuming the life cycle is a Markov chain with rewards (Caswell, 2011; van Daalen and Caswell, 2017, 2020). Specifically, we defined the life-cycle transition as a multistate model indexed by ages *x* = 1, …, *ω* and years *t* = 1, …, *g*; for details of such models, see Caswell et al. (2018). Thus, the population projection matrix is

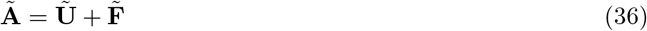

where the survival matrix 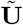 and fertility matrix 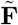 each having year-within-age block structure. We constructed the matrices 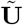 and 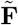 from vital rates specific to age and year, capturing both types of transitions. First, we defined the *ω × ω* matrix **U**_*i*_ that describes the age transition for year *i* having survival probabilities on the subdiagonal and zeros elsewhere. We incorporated the kin-mediated feedback by reparameterising the survival matrix using (33) and then solving the nonlinear equations (20) using nleqslv (Hasselman and Carnell, 2026). Hence, we obtained the solved survival matrix 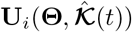. However, for notational simplicity, we do not mention the parameters in parentheses in the subsequent sections. Furthermore, we defined the *g × g* matrix ***D***_*j*_ to describe the transition among years for individuals in age *j*

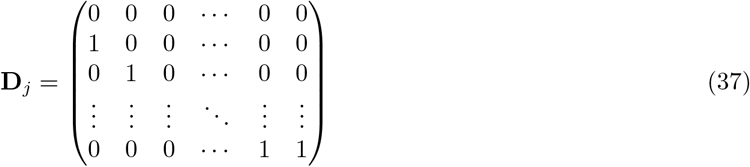

where year sequences are deterministic but aperiodic, moving through a specified sequence of matrices, *i*.*e*., the temporal sequence of matrices from 1950 to 2023, we observed in the data. Then, using **U**_*i*_ and **D**_*j*_ we constructed block-structured projection matrix as

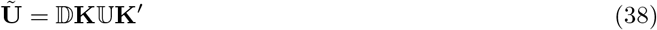

where **K**_*g,ω*_ is a vec permutaion matrix that satisfies vec(**X**^*′*^) = **K**_*g,ω*_vec**X**. The *gω × gω* block diagonal matrices U and D are defined as

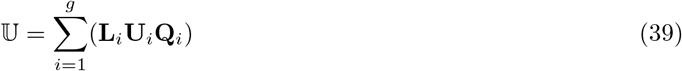

where the block construction matrices **L**_*i*_ and **Q**_*i*_ are given by

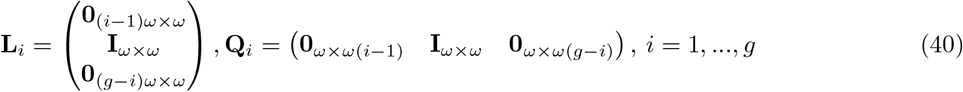

To compute the mean LRO, we expressed the life cycle as a Markov chain with rewards. Specifically, we defined the life cycle transition as a Markov chain transition matrix

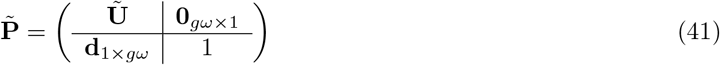

where the final state, in the last row of 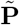 is the absorbing state of death. Reproduction is described as a *reward* that individuals accumulate as they survive and move among the ages of their life cycle (Caswell, 2011; van Daalen and Caswell, 2017, 2020). The reproductive rewards associated with each transition age given by matrix **R**_*k*_ where the entry of **R**_*k*_ is the *k*th moment of the reproductive output associated with each transition. The matrix 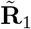 is a function of

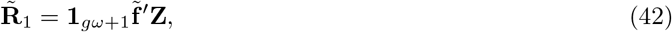

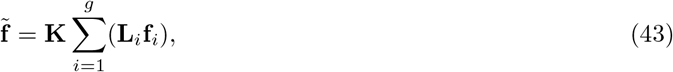

where **f**_*i*_ is the fertility vector that appears on the first row of **A**_*i*_ and **Z** = (**I** | **0**) is a matrix that cleaves off the absorbing state, because the dead do not reproduce (Caswell, 2011; van Daalen and Caswell, 2017, 2020). Thereafter, we computed the first moment of LRO as

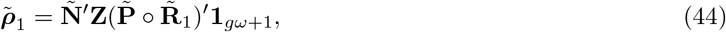

where

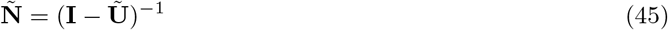

is the multistate fundamental matrix (van Daalen and Caswell, 2020). Note that the LRO accounts for the nonlinear feedback between the kinship network and survival rates as we used the solved survival matrix 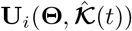 as the input for LRO computation. Furthermore, the age *×* time model in (41) can go up to age 74 as we have 74 Leslie matrices for the period 1950-2023. However, this censoring at age 74 does not affect the calculation of LRO, as reproduction finishes much before that age.

### LRO Sensitivity

To test our hypotheses (H1 and H2) regarding the sensitivity of LRO to kin dependence, we computed the derivative of the first moment of LRO 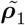 to slope coefficients ***γ***. As our parameter set of interest is the slope coefficient ***γ*** of kins *K* we computed the sensitivity of mean LRO as

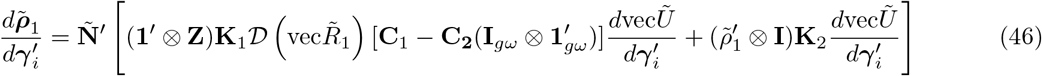

where

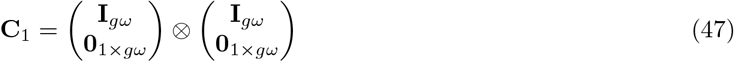

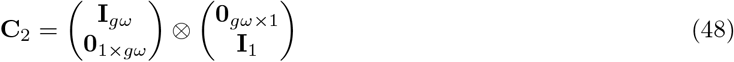

and **K**_1_ = **K**_(*gω*=1),(*gω*+1)_, and **K**_2_ = **K**_*gω,gω*_ are the vec-permutation matrices (van Daalen and Caswell, 2017, 2020). Here we obtained the derivative of *γ*_*i*_ to the block-structured survival matrix 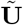 as

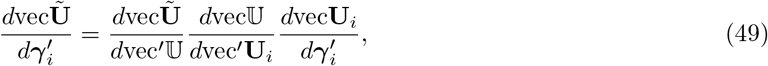

or

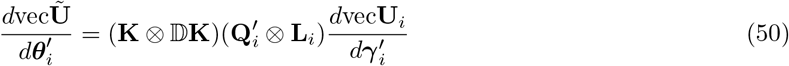

following the protocol proposed by van Daalen and Caswell (2017, 2020).

## Results

We find strong support for our hypothesis H1a, regarding the greater dependence of child survival on the mother’s presence being under positive selective pressure. Specifically, our results show that the sensitivity of the stochastic growth rate to the slope coefficient of the probability of having a living mother on child mortality 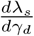 is negative across all three examined population types (panel A in Figure 2). Hence, increasing the negative slope coefficient *γ*_*d*_ = −0.947, *i*.*e*., taking it closer to zero, decreases the stochastic growth rate *λ*_*s*_ for each of the three types of population. In other words, reducing the dependence of child mortality on the mother’s presence would be under negative selection. Conversely, increasing the absolute magnitude of the negative slope coefficient for mothers’ presence *γ*_*d*_ would increase the stochastic growth rate. The sensitivity is highest in the *type-1* population and slightly lower in the *type-2* population (panel A in Figure 2). However, the sensitivity for the *type-3* population, which is in the latest stage of demographic transition and has low child mortality, is considerably lower than for the other two population types.

**Figure 2:**
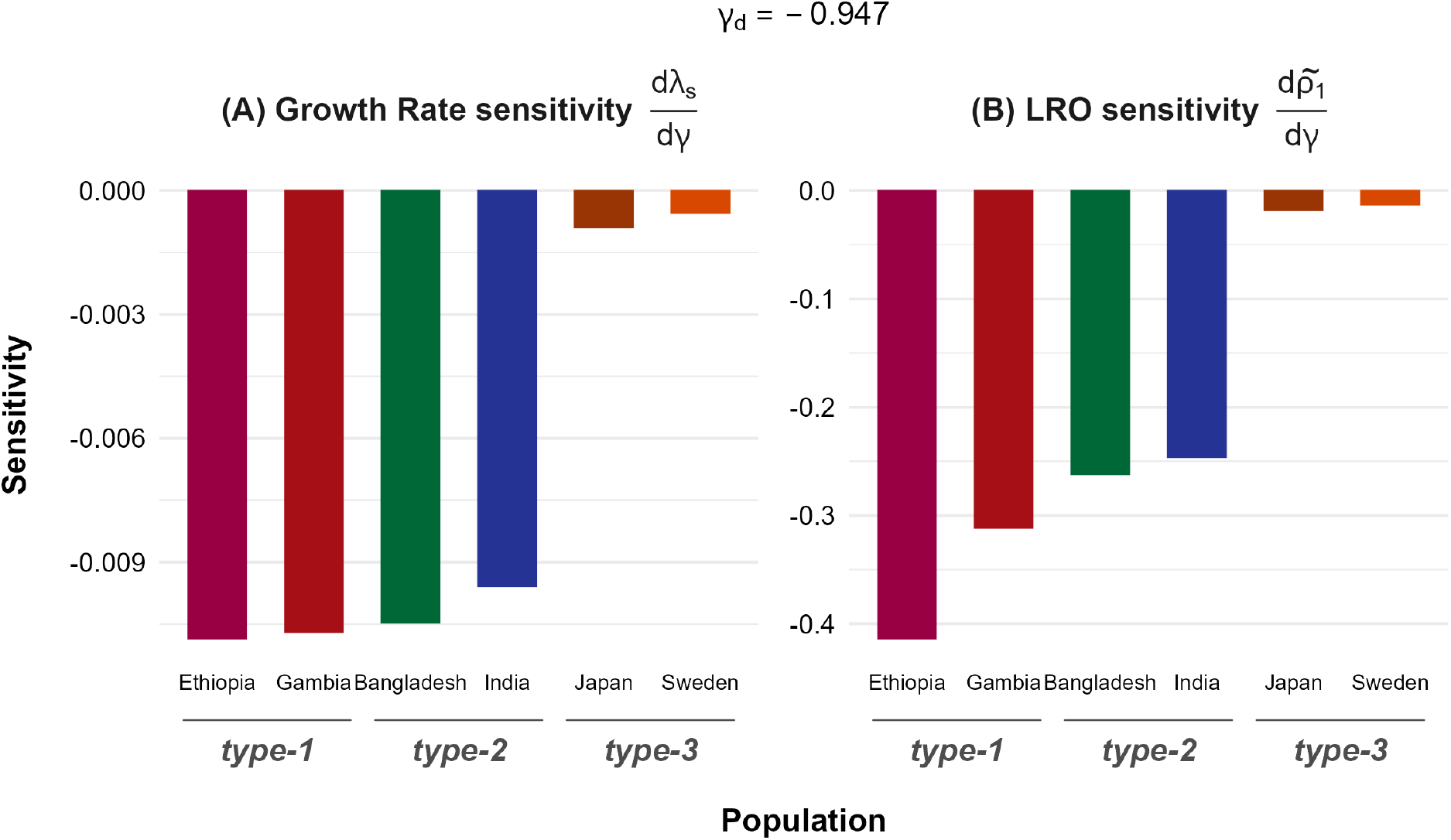
Negative sensitivity of both the stochastic growth rate *λ*_*s*_ and mean lifetime reproductive output (LRO) 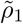 to the slope of the probability of having a living mother on child mortality *γ*_*g*_ represents a negative selection pressure on *γ*_*g*_. Panel (A) shows the sensitivity of the stochastic growth rate 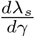 and panel (B) shows the sensitivity of the mean LRO 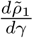 to the slope coefficient *γ*_*d*_. Here, separate bars illustrate the sensitivity for each country representing one of the three population types. The value of the slope coefficient *γ*_*d*_ is displayed at the top of the figure.

In line with our expectations, the negative sensitivity of mean lifetime reproductive output (LRO) also supports our hypothesis H1b regarding the dependence of child mortality on maternal presence. We find that the sensitivity of mean LRO to perturbation in the slope coefficient of probability of having a living mother on child survival 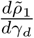 is negative across all population types. The negative sensitivity indicates that an increase in the negative value of *γ*_*d*_, *i*.*e*., reducing the negative slope toward zero, would decrease the stochastic growth rate. Hence, reducing the dependence of child mortality on the availability of a living mother would reduce the mean LRO. Contrary to the stochastic growth rate, the sensitivity of mean LRO for *type-1* population is noticeably higher than the *type-2* population (panel A in Figure 2). Moreover, the sensitivity of the mean LRO is much larger in magnitude than that of the stochastic growth rate. Similar to the former case, the sensitivity of the mean LRO, the slope coefficient of having a living mother in *type-3* population, is considerably lower than the other two population types.

Our results support hypothesis H1b, which posits that the dependence of child mortality on grandmother’s presence is under positive selection pressure. We tested this hypothesis by examining whether the sensitivity of the stochastic growth rate to the slope coefficient of the probability of having a living grandmother on child mortality 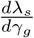 is negative across all population types. As expected, we found negative sensitivities across all population types (see panel A of Figure 3). A corollary to this finding suggests that an increase in the absolute value of the negative slope *γ*_*g*_ (*i*.*e*., a decrease in the negative slope) would increase the stochastic growth rate. Hence, selection favours greater dependence of child mortality on the presence of a living grandmother. However, contrary to the sensitivity to grandmothers’ effect, *type-2* population shows higher sensitivity to the slope coefficient of mother’s presence *γ*_*g*_ than that of the *type-1* population; although the difference is small. In agreement with the sensitivity to the mother’s effect, sensitivity to the slope coefficient of the grandmother’s presence *γ*_*g*_ is also drastically lower in *type-3* population, where the child mortality is very low ~ 0.002, compared to the other types. Overall, the sensitivity of the stochastic growth rate to *γ*_*g*_ is lower than that of the mother’s slope coefficient, indicating that perturbing the slope of the mother’s presence on child mortality would bring larger fluctuations in the stochastic growth rate than that of the grandmother’s presence.

**Figure 3:**
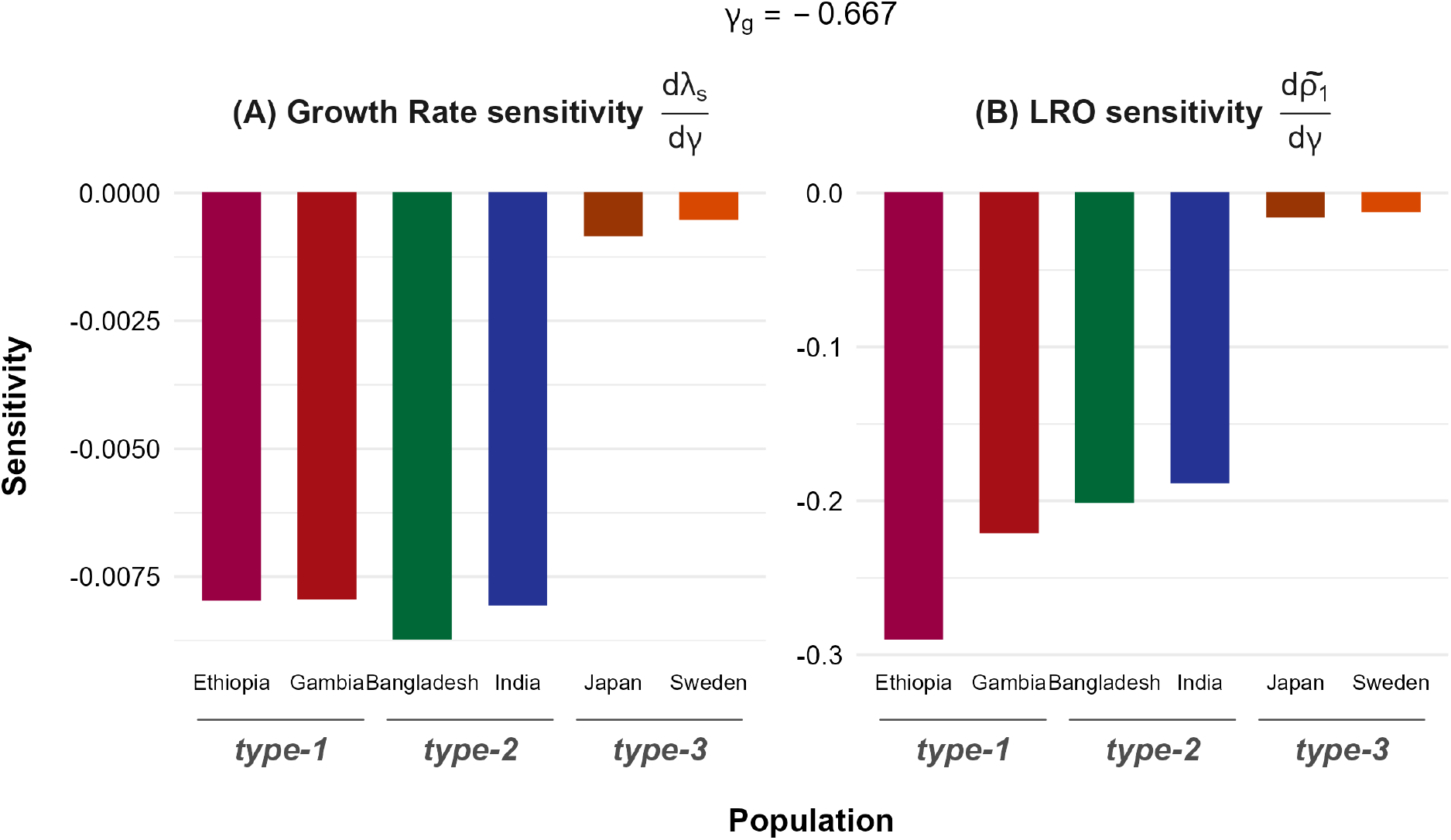
Negative sensitivity of both stochastic growth rate *λ*_*s*_ and mean lifetime reproductive output (LRO) 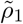 to the marginal effect of grandmothers *γ*_*g*_ indicates a negative selection pressure. Panel (A) shows the sensitivity of the stochastic growth rate to the slope coefficient of the probability of having a living grandmother on child mortality 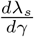 with bars for each type of population. Panel (B) illustrates the sensitivity of lifetime reproductive output to the perturbation in the slope coefficient for the effect of having a living grandmother on child mortality. Here, the bars display sensitivities for countries representing each population type. The slope coefficient *γ*_*g*_ value is shown at the top of the figure.

We find similar results from the perturbation analysis of the mean LRO with respect to the slope coefficient of grandmother’s presence. Specifically, our result shows that the sensitivity of the mean LRO to the slope coefficient for grandmother’s presence, 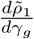, is negative across all population types. In other words, increasing the negative slope coefficient *γ*_*g*_ = −0.667, *i*.*e*, taking it closer to zero, would decrease the mean LRO across all population types. Hence, reducing the dependence of child mortality on the grandmother’s presence would lower the focal individual’s mean LRO. However, the sensitivity in *type-1* population is highest among the three population types, with a noticeable difference from that of *type-2* population (panel B in Figure 3). In the population *type-3*, which is the advanced level of demographic transition with the lowest child mortality, the sensitivity of the mean LRO to the slope of grandmother’s presence on child mortality is also the lowest among the three population types. The sensitivity of the mean LRO is also much larger than the stochastic growth rate’s sensitivity to the slope of grandmother’s presence on child mortality. The highest LRO sensitivity observed across the three population types is ~ 0.3, whereas the stochastic growth rate sensitivity is only ~0.01.

In support of our hypothesis H1c, we find that selection favours a greater dependence of child mortality on the presence of a living older sister (*>* 10 years). Our result shows that the sensitivity of the stochastic growth rate to the slope coefficient of the probability of having a living older sister age *>* 10 years on child mortality 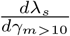 is negative across all three examined population types. Increasing the slope coefficient *γ*_*m>*10_ = −0.239, *i*.*e*., taking it closer to zero, would reduce the stochastic growth rate. The opposite is true, as increasing the value of |*γ*_*m>*10_|, *i*.*e*., decreasing the negative slope, would lead to an increasing stochastic growth rate. Hence, increasing dependence of child mortality on the presence of an older sister of age *>* 10 would be under positive selective pressure. However, the magnitude of the sensitivity varies considerably across the population types. *Type-1* populations show the highest sensitivity ~0.003, followed by the *type-2* population with the magnitude ~0.0015 (panel A in Figure 4). Interestingly, the sensitivity in the *type-3* population is very close to zero, indicating that perturbations in the slope coefficient of an older sister’s (*>* 10 years) presence on child mortality do not translate into a noticeable fluctuation in the stochastic growth rate. It is also important to note that the absolute value of the stochastic growth-rate sensitivity for older sisters (*>* 10 years) is much smaller than that for the mother or grandmother.

**Figure 4:**
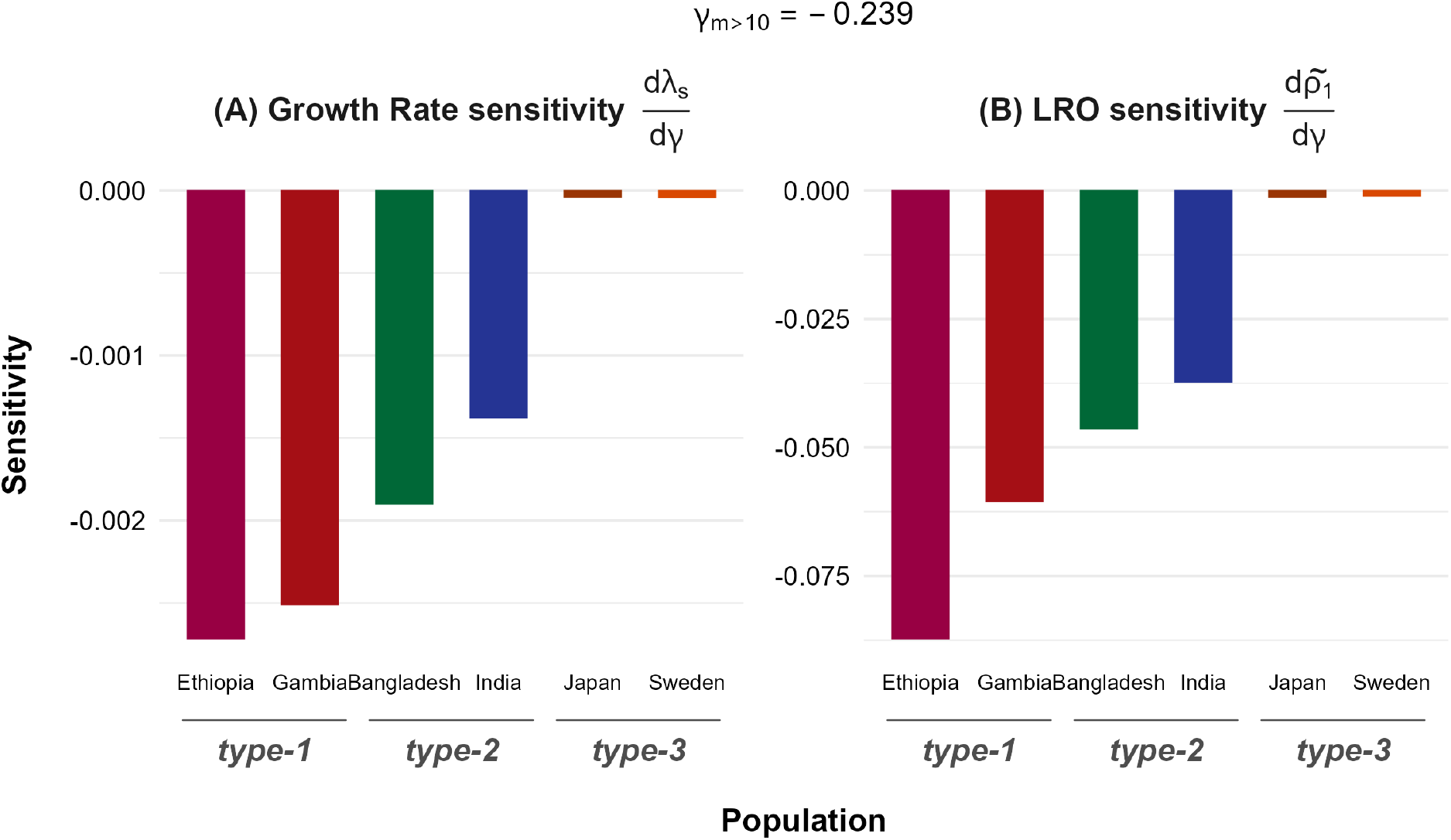
Negative sensitivity of stochastic growth rate *λ*_*s*_ and mean lifetime reproductive output (LRO) 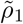 to the marginal effect of having a living older sister (*>* 10 years) on child mortality indicates a negative selection pressure. Panel (A) illustrates the sensitivity of growth rate 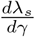, and panel (B) shows the sensitivity of mean lifetime reproductive output 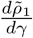 to the slope coefficient of the probability of having a living older sister aged *>* 10 years on child mortality *γ*_*m>*10_. The bars represent the sensitivities for each population type. The magnitude of the slope coefficient *γ*_*>*10_ is displayed at the top of the figure.

The mean LRO sensitivity with respect to perturbations in the slope coefficient *γ*_*m>*10_ also supports our hypothesis H1c regarding older sisters (*>* 10 years). We find that the sensitivity of the mean LRO to the slope coefficient for the older sister’s (*>* 10 years) presence on child mortality, 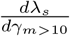, is negative across all three population types examined. As with mothers and grandmothers, sensitivity in the *type-1* population is highest, but the difference from that in the *type-2* population is more pronounced for older sisters (*>* 10 years). The sensitivity in the *type-3* population is also closer to zero for the mean LRO. Moreover, the highest absolute value of mean LRO sensitivity 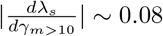 (panel B in Figure 4) is considerably lower than that of the mother (~0.4) and grandmother’s (~0.3). Thus, perturbing the slope coefficient of the probability of having a living older sister (*>* 10 years) on child mortality does not produce the same fluctuation as it does when the same is done for the mother or grandmother.

We find strong support for the hypothesis H2 that the association of child mortality with the presence of an older sister aged ≤10 is under negative selection. Specifically, our result shows that the sensitivity of the stochastic growth rate to the slope of the probability of having a living older sister (≤ 10 years) on child mortality 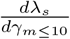 is negative across the three population types. As the slope coefficient for the probability of having an older sister (≤ 10 years) on child mortality *γ*_*m*≤10_ = 0.21 is positive, the negative sensitivity indicates that an increase in slope coefficient *γ*_*m*≤10_ would lead to a decrease in the stochastic growth rate across all population types. Moreover, we find that the sensitivity in the *type-1* population is highest with a magnitude around ~0.005, followed by that in the *type-2* population ~0.004 (panel A in Figure 5). Similar to the previous kin, the stochastic growth rate sensitivity in the *type-3* population is very small. Overall, the highest absolute sensitivity is ~ 0.006, slightly smaller than that of the mother and grandmother but considerably larger than that of the older sisters (*>* 10 years).

**Figure 5:**
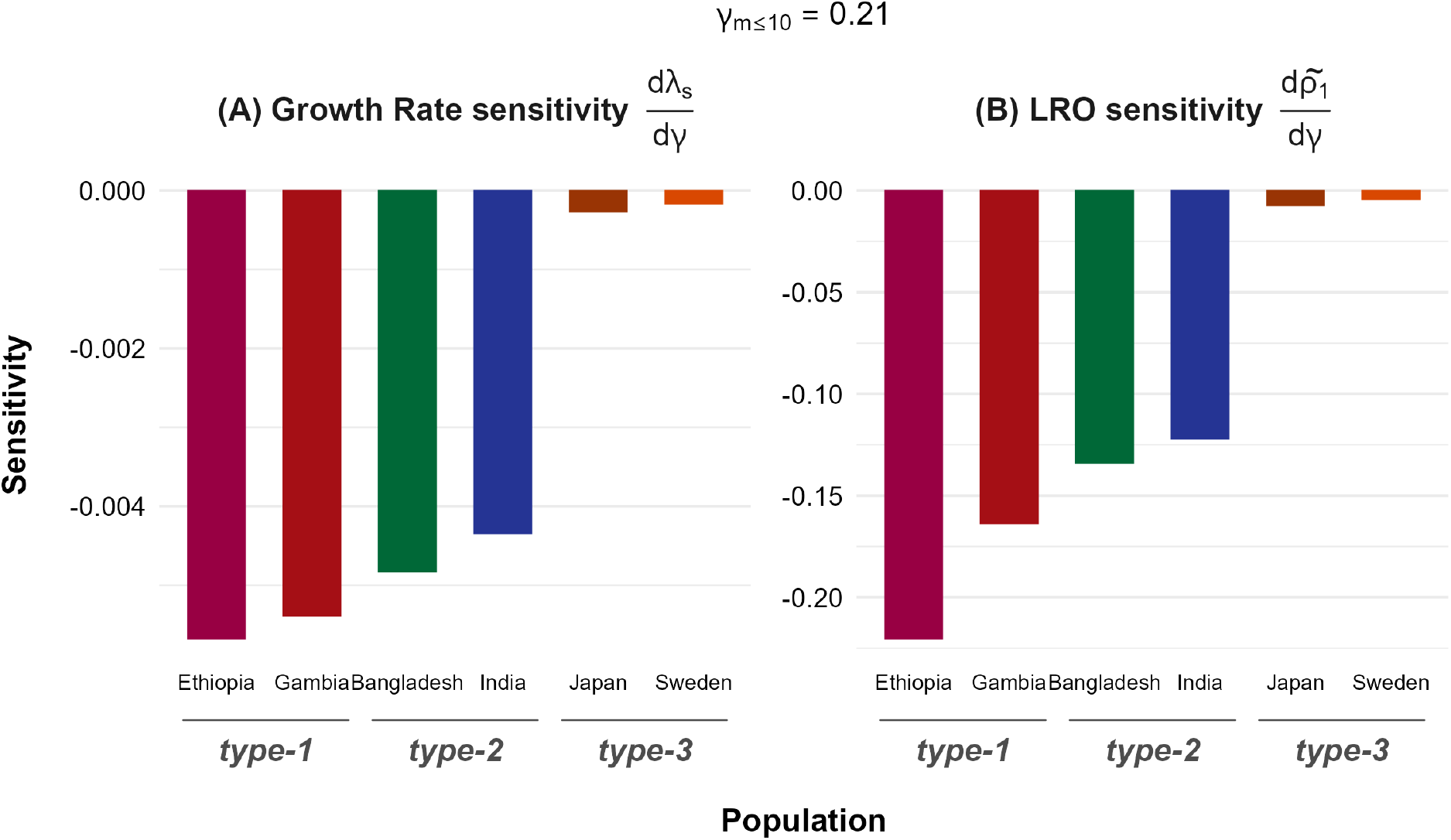
Negative sensitivity of stochastic growth rate *λ*_*s*_ and mean lifetime reproductive output (LRO) 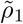 to the positive marginal effect of having a living older sister ≤10 years on child mortality indicates a negative selection pressure on the effect. Panel (A) illustrates the sensitivity of the stochastic growth rate 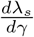 and panel (B) shows the sensitivity to the mean lifetime reproductive output 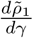 to the slope coefficient of the probability of having a living older sister aged ≤10 years on the child mortality *γ*_≤10_. Each bar illustrates the sensitivities for each country representing a population type shown on the x-axis. The value of the slope coefficient *γ*_≤10_ is displayed at the top of the figure.

The mean LRO sensitivity with respect to perturbation in *γ*_*m*≤10_ also supports the hypothesis H2. We find that the sensitivity of the mean LRO to the slope coefficient *γ*_*m*≤10_, *i*.*e*., 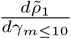 is negative for all the population types. The results show considerable variation in the magnitude of sensitivity across population types. Specifically, the sensitivity in *type-1* population is ~ −0.2, followed by *type-2* population with sensitivity ~ −0.15, and finally the *type-3* population shows a sensitivity very close to zero (panel B in Figure 5). Interestingly, the absolute value of the sensitivity between the stochastic growth rate is considerably lower than that of the mean LRO, indicating that the perturbation in slope coefficient *γ*_*m*≤10_ leads to a much larger fluctuation in the mean LRO than in the stochastic growth. However, caution is warranted when comparing the sensitivity of the stochastic growth rate and the mean LRO, as both metrics are computed over different timescales and under different environmental conditions. We computed the stochastic growth rate under an iid environment over 1000 time steps, while the mean LRO was computed for a deterministic but aperiodic environment over the observed 74-year sequence.

## Discussion

Examining the coevolution of kin-derived sociality and senescence can provide important insights into the extension of human longevity and the role of kin selection. However, studies examining selection on kin-derived sociality do not explicitly model the dynamic feedback between the kinship network and the focal individual’s life history, which leads to the longevity extension (Carey and Judge, 2001). Cyrus Chu and Lee (2006) computed the sensitivity of the growth rate to an intergenerational (resource/energy) transfer, without explicitly considering the kinship network, whereas Pavard and Branger (2012) computed the sensitivity to age- and kin-status-specific survival rates rather than to the slope of kin availability on mortality. Although Croll and Caswell (2025) modelled the nonlinear feedback between the kinship network and the life history of the focal individual, they compared population growth across scenarios rather than computing the sensitivity of the growth rate to the kin coefficient. To address this gap, we computed the derivative of the stochastic growth rate with respect to a perturbation in the slope coefficient of kin availability on child mortality while accounting for the kinship-mortality feedback. Our results show a negative growth sensitivity for the negative slope of mother, grandmother, and older sister (*>* 10 years) on child mortality across all three examined population types. The findings can be interpreted as indicating that reducing the dependence of child mortality on the presence of the abovementioned kin (by moving the negative slope towards zero) would be under negative selection pressure, or, conversely, that selection favours a greater kin dependence of child mortality. Interestingly, our finding that selection on kin dependence of juvenile survival is strongest in the *type-1* population with the highest child mortality and progressively weakens under lower-mortality regimes is consistent with the pattern observed by Pavard and Branger (2012). Moreover, existing studies do not discuss the limits of the self-reinforcing mechanism between the kin effect and survival (Bourke, 2007); our result suggests that progress in reducing baseline mortality might limit this mechanism in human populations.

We show the sensitivity of individual fitness to the kin dependence via mean lifetime reproductive output (LRO). The first moment of LRO is considered a critical component of fitness in evolutionary biology (Caswell, 2011), and its sensitivity to perturbations in the slope coefficient of kin presence can provide important insights into individual fitness (Tuljapurkar et al., 2009; van Daalen and Caswell, 2017, 2020). Similar to the stochastic growth rate, increasing the negative slope coefficient, *i*.*e*., averting the negative relationship of the presence of mother, grandmother, and an older sister (*>* 10 years) with child mortality, would reduce the lifetime reproductive output. As with the stochastic growth rate, averting the negative association of maternal, grandmaternal, and older-sister (*>* 10 years) presence with child mortality increases the mean LRO, with the sensitivity magnitude largest in the high-mortality *type-1* population and near zero in the low-mortality *type-3* population. This pattern closely matches van Daalen and Caswell (2017) finding, across a comparable range of human demographic regimes, that the sensitivity of mean LRO to age-specific mortality is largest in historically high-mortality, high-fertility populations (the Ache, Hadza, and Hutterites) and smallest in low-mortality modern populations (the Netherlands, Sweden, and Japan), as high fertility magnifies the fitness consequences of a mortality change while low baseline mortality leaves little room for a marginal change to matter. Our result extends this pattern from mortality in general to the kin dependence (lower-level parameter) of mortality specifically, distinguishing the marginal fitness contribution of kin presence from that of baseline mortality. Although it might be tempting to infer positive selection pressure from negative sensitivity, the measure is not computed on a long time scale, as in the stochastic growth rate and therefore does not provide a signal of selection.

The positive association of competing kin with child mortality is under negative selection pressure. As the competitive and cooperative effects of sibling interaction have been well documented across taxa, namely insects (Andersson, 1984), birds (Koenig and Dickinson, 2004), and mammals (Solomon and French, 1997), we also examined the fitness impact of an elder sister below the age of maturity (*<* 10 years) as a competing kin alongside helping sisters (*>* 10 years). We find that increasing the positive association of elder sister (*<* 10 years) with child mortality, potentially due to competition over resources, such as (grand) parental care, would reduce the stochastic growth rate. Empirical studies suggest that higher-order birth and a shorter birth interval between siblings, i.e., an elder sister with a smaller age gap to the focal, increase the focal’s mortality risk (Riswick and Hsieh, 2020; Thomas et al., 2015; Whitworth and Stephenson, 2002). However, the strength of sibling interaction is influenced by factors at the individual level (e.g., birth order), the family level (e.g., socioeconomic conditions, mother’s education), and the population level (e.g., life history, stage of demographic transition) (Nitsch et al., 2013; Whitworth and Stephenson, 2002). In line with this argument, we find that perturbation of the slope coefficient of the older sister’s (*<* 10 years) on child mortality leads to a relatively larger fluctuation in stochastic growth rate for the *type-1* population at an early stage of demographic transition, but almost zero fluctuation for the *type-3* population in a very low mortality regime. Moreover, the existing literature on the demographic impact of sibling interaction is limited to short-term measures, such as physical growth and survival to age at maturity (but see Nitsch et al., 2013). In this study, we examined the sensitivity of mean lifetime reproductive output to stochastic growth rates and showed that increasing the negative relationship of competing sisters with the focal child’s survival reduces the expected LRO, thereby diminishing individual fitness. Our result aligns with Nitsch et al.’s (2013) result that the negative influence of elder sisters on female reproductive success reduces lifetime fitness; however, their result applies to all elder sisters, not just those under 10 years old.

We provide a framework for examining the coevolution of life history and sociality, as well as the natural selection acting on sociality. Our framework advances the discipline in three key ways. First, we explicitly model the nonlinear feedback between the life history and the kinship network of the focal individual, incorporating environmental stochasticity into the feedback loop. One of the first formal models to examine the coevolution of mortality and sociality was provided by Cyrus Chu and Lee (2006), who quantified selection on a transfer from age *j* to age *i* as the derivative of the long-run growth rate with respect to the transfer, showing that reducing mortality and rising transfers reinforce each other. However, the model does not consider the (genetic) relatedness between the provider and the recipient of the intergenerational transfer, and a transfer can evolve only if the interactants are genetically related (Hamilton, 1964). Pavard and Branger (2012) incorporated relatedness directly, using a matrix population model classified by age and (grand) mothers’ survival status and then computed the sensitivity of the long-term growth rate to survival rates to show how (grand) maternal care strengthens selection on survival. Although the model neatly incorporates the direct effect of closely related kin on an individual, it does not account for how these effects percolate through the kin network. Croll and Caswell (2025) bridge this gap by providing an advanced model that accounts for nonlinear feedback between the entire kinship network and the life history of the focal individual, which we extend by incorporating environmental stochasticity in the feedback.

Second, we develop a Bayesian hierarchical model to express the focal individual’s mortality as a function of kin survival. In the absence of a parametric mortality model to incorporate the kin dependence, previous studies have quantified mortality either by borrowing the relative risk of mortality in the absence of a particular kin (Pavard and Branger, 2012) or borrowing point estimates of regression coefficients from published studies (e.g., Parker et al., 2021) as a multiplicative adjustment factor to survival rates (Croll and Caswell, 2025). In both cases, the borrowed parameter is treated as a known constant, with no mechanism to propagate the sampling uncertainty of the original estimate into uncertainty around the estimated mortality (or survival). We provide a flexible Bayesian hierarchical model that expresses focal mortality as an explicit function of kin availability, with the slope coefficient of the kin presence modelled with an informative prior derived from published studies. Our model allows us to formally incorporate prior knowledge from the same body of literature that previous studies drew single-point estimates from, while yielding a full posterior distribution of the slope of kin presence on mortality. Consequently, we can estimate mortality by propagating uncertainty from published studies on the regression coefficient into uncertainty around our estimates.

Finally, we examine selection on sociality by computing the derivative of the stochastic growth rate with respect to the kin dependence. Our analysis is a stochastic analogue of Cyrus Chu and Lee (2006) deterministic sensitivity of the growth rate to intergenerational transfer and an extension of Pavard and Branger’s (2012) sensitivity of the growth rate to age- and kin-survival-status-specific survival rates to a lower-level parameter. Additionally, we estimated the mean LRO as a function of the kinship network and computed its sensitivity to the kin dependence. It is important to note that van Daalen and Caswell (2017, 2020) provided formulations for sensitivity analysis of mean LRO across variable environments, and van Daalen et al. (2022) presented lifetime reproductive output as a function of the mother’s mean age. Additionally, empirical studies provide ample evidence of the effect of kin presence on human fertility and child survival (Sear et al., 2003; Sear, 2018; Snopkowski and Sear, 2016, 2013). To the best of our knowledge, this is the first study to present the sensitivity of the mean LRO to perturbations in the slope coefficient of kin presence on survival rates, while accounting for feedback between the rates and the kinship network.

Some key limitations of our study demand caution in making inferences from the results presented here. First, we do not account for demographic stochasticity in constructing the kinship network; however, demographic stochasticity can introduce significant fluctuations in kinship networks that our model fails to capture (Caswell, 2024). Second, our matrix population models are female-dominant; therefore, our results fail to capture kin-mediated population dynamics in overall human populations. Third, we assume a constant sex ratio at birth of 1.05 to make the fertility rates sex-specific, which does not reflect the actual sex composition of births. Fourth, we derive the informative priors for the slope coefficients of the probability of having a specific kin on child mortality from a very limited number of studies based on a specific population (Jamison et al., 2002; Sear and Mace, 2008, 2022), which makes the priors restrictive. Moreover, we do not perform a prior sensitivity analysis to validate the priors on the slope coefficients in ***γ***; therefore, the posterior estimates of these coefficients should be interpreted with caution. Fifth, we present the sensitivity results only with respect to the median of the posterior estimates of the slope coefficients, as the analysis for samples of ***γ*** posterior is computationally very expensive. Hence, we do not show the uncertainty in the sensitivity translated from the uncertainty in the slope coefficient, a strength of Bayesian estimation. Finally, in our study, we examine the association between specific kin and child mortality and do not consider the effect of having a specific kin on the focal’s fertility, which can have a significant impact on asymptotic population dynamics.

Our framework can be extended to address the aforementioned limitations and applied across a range of contexts. Future research in this area could focus on developing formulations that incorporate the potential impact of demographic stochasticity on the nonlinear feedback between the kinship network and vital rates. Another important extension of the framework is to expand it to two-sex and/or multistate models, so that it can better capture the dynamics of species for which (developmental) stage transitions are more important than age transitions. Additionally, in our formulation, we assume the environmental states are independent and identically distributed, with no environmental autocorrelation; this assumption can be extended to include temporal autocorrelation via a finite-state Markov chain transition matrix for the environment (Caswell, 2001). Moreover, the informed prior on the slope coefficient of kin availability, ***γ***, can be improved by conducting a meta-analysis of studies examining kin influence on child survival among humans and by performing a prior sensitivity analysis. If computational resources permit, the results can be presented with an uncertainty interval around the sensitivity estimates, reflecting the uncertainty in the posterior distributions of the slope coefficients. Additionally, incorporating kin feedback into fertility rates by developing another Bayesian model for fertility would be a future direction for our research. Finally, an important application of our framework in ecology and evolutionary biology would be to examine how non-linear feedback between the kinship network and vital rates affects demographic buffering against stochastic environments (Hilde et al., 2020; Pfister, 1998).

The examination of the coevolution of kin interaction and ageing needs to incorporate nonlinear feedback between the kinship network and vital rates. Our work provides a quantitative framework for examining the sensitivity of the stochastic growth rate to kin effects on survival, accounting for the self-reinforcing feedback between kin and survival rates in stochastic environments. Therefore, we can infer selection pressure on kin interactions and how they coevolve with life histories. Here, we have quantified selection pressures on kin interactions in human populations across different demographic regimes and shown how individual fitness may fluctuate in response to changes in kin interactions. Our study contributes to the existing literature on the coevolution of sociality and ageing by providing a quantitative framework to examine the sensitivity of stochastic growth rate to kin interaction while accounting for the nonlinear feedback loop.

## Appendix

### Prior Specification

To model the slope of the kin covariates, we defined informative priors on the slope coefficients. We obtained the slope coefficients and standard errors for the effects of living with a mother, a grandmother, and older sisters from the studies by Jamison et al. (2002) and Sear and Mace (2008, 2022). Then, we pooled the slope estimates and standard error for each covariate from these studies using the inverse-variance weighting scheme

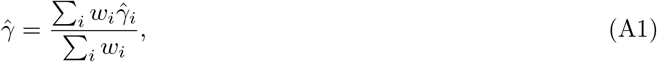

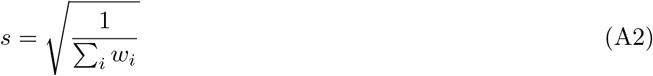

where the weight is 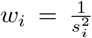 for each study *i*. Finally, we define the prior on slope coefficient for each covariate in ***K*** as 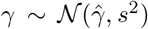. The informative prior on the ***γ*** slope coefficients defined based on the published studies were as follows:

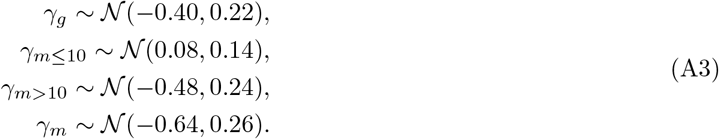

It is worth noting that we have obtained estimates of slope coefficients from three studies, as studies examining the effect of kinship on child mortality with large sample sizes are scarce. However, a detailed meta-analysis of studies investigating the kin effect on child mortality and the construction of a more robust prior are exciting topics for future research.

To model the sharp spikes in mortality, we defined a regularised horseshoe prior on the parameter *δ*_*tl*_. Following Piironen and Vehtari (2017), we expressed the prior on *δ*_*tl*_ as

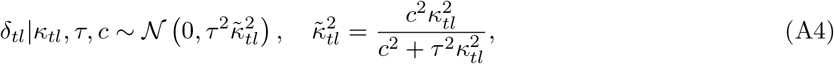

where the global hyperparameter *τ* shrinks all the *δ*_*tl*_ towards zero, while the heavy-tailed half-cauchy prior on local hyperparameter *κ*_*tl*_ ~ *C*^+^(0, 1) allows some *δ*_*xl*_ to escape the shrinkage when data strongly suggests it. The parameter 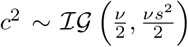 ensures that it not only shrinks small *δ*_tl_ as original horseshoe prior (Carvalho et al., 2010) but also regularises the largest *δ*_*tl*_ with a Student-*t*_*?*_ (0, *s*^2^) slab. Finally, following the recommendation of Piironen and Vehtari (2017), we defined the prior for the global shrinkage parameter *τ* as 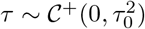, where

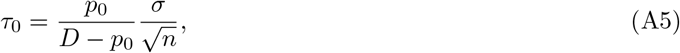

where *p*_0_ is our prior guess for number of relevant variables for *δ*_*tl*_, *D* is the number of variables, *n* is the number of data points and *σ* is the noise level. The regularised horseshoe prior on the random country-year fluctuation parameter *δ*_*tl*_ ensures that it deviates the mortality curve to fit the sharp mortality spike when data suggests it otherwise shrinks towards zero. Furthermore, we defined priors on the hyperparameters of the regularised horseshoe prior as

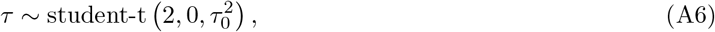

where

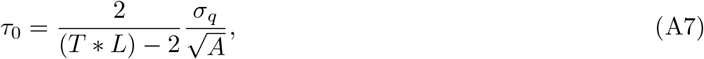

with *T* = 74, *L* = 5, and *A* = 5. The standard deviation for the mortality is defined as

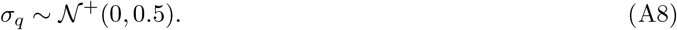

Priors for the remaining parameters were defined as

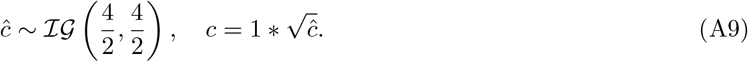

To complete the prior specification, we defined a weakly informative prior on the remaining parameters. Specifically, to ensure that no inforamtion on the level and trend of the spline term Φ_*xl*_(*t*) was shared across ages and countries, we defined the priors on the age-country specific level and slope parameters as *α*_*xl*,0_ ~ *N* (0, 5) and *α*_*xl*,1_ ~ *N* (0, 1), respectively. On the other hand, we allow information sharing in spline coefficients across age and country through a hierarchical prior on the variance parameter for the second-order difference in spline coefficients,

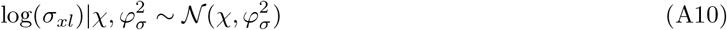

(Alkema and New, 2014). Finally, we defined priors on the hyperparameters

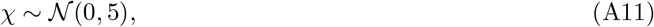

and

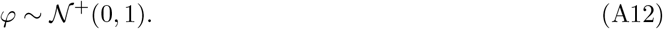

### Parameter Estimation

To perform the inference on the parameters in (23), we sampled the posteriors using the Hamilton Monte Carlo algorithm (HMC, hereafter). Specifically, we employed the No-U-Turn Sampler (NUTS; Hoffman and Gelman, 2014), a variant of HMC, via the probabilistic modelling software Stan, accessed through the cmdstanr interface in R (Gabry et al., 2025). Furthermore, we assessed the convergence of the parameters using the measures of split-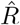 as well as tail and bulk effective sample size (ESS, hereafter). The split-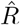 measures the total variance relative to within-chain variance, indicating how much the posterior samples vary within each chain. Thus, the posterior samples should be used only if the split-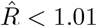, suggesting that the total variance is similar to within-chain variance, *i*.*e*., all the chains are exploring the same distribution of the parameters. Additionally, we check the bulk and tail ESS, which refer to the ESS of the rank-normalised draws and the minimum of the ESS of the 5% and 95% quantiles. Following the recommendation of Vehtari et al. (2021), we use the posterior samples only if both the ESS measures exceed 100 times the number of chains.

We fit the model using four separate chains with 4000 iterations, of which 2000 iterations were used as warm-ups. The measure of split-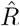 were less than 1.01 for all the parameters, and both measures of ESS were more than 400 for each parameter, indicating convergence for all models and parameters.

### Model Checks and Evaluation

To validate the Bayesian model, we conducted posterior predictive checks, following REF. Specifically, we simulated replicate data for the **q**_rep_ using the posterior predicitive distribution

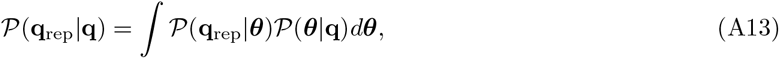

where **q** are the observed mortality and ***θ*** are the parameters in (23). Then, we compared the densities of the observed and replicated data to check whether the model-generated data approximated the observed mortality data. Additionally, we computed test statistics for the mean and standard deviation, *T* [**q, *θ***], to compare replicated and observed mortality. We check whether the test statistics from the observed mortality lie near the centre of the histograms constructed from the test statistics of the replicated mortality data to assess model fit. Furthermore, we prepared trace plots of the slope coefficients in ***γ*** to show their convergence. Finally, we evaluated the posterior of the mortality using the mean absolute error and the root mean squared error. We obtained a mean absolute error of 0.0004 and a root-mean-square error (RMSE) of 0.0012.

To illustrate the model fit, we provide figures of the density overlay (see Figure A1 and Figure A2), residuals (Figure A3), and the fit of replicated mortality derived from model posteriors to the observed mortality (Figure A4). We also provide histograms of the test statistics (see Figure A5 and Figure A6) and trace plots of the ***γ*** slope coefficients (Figure A7).

**Figure A1:**
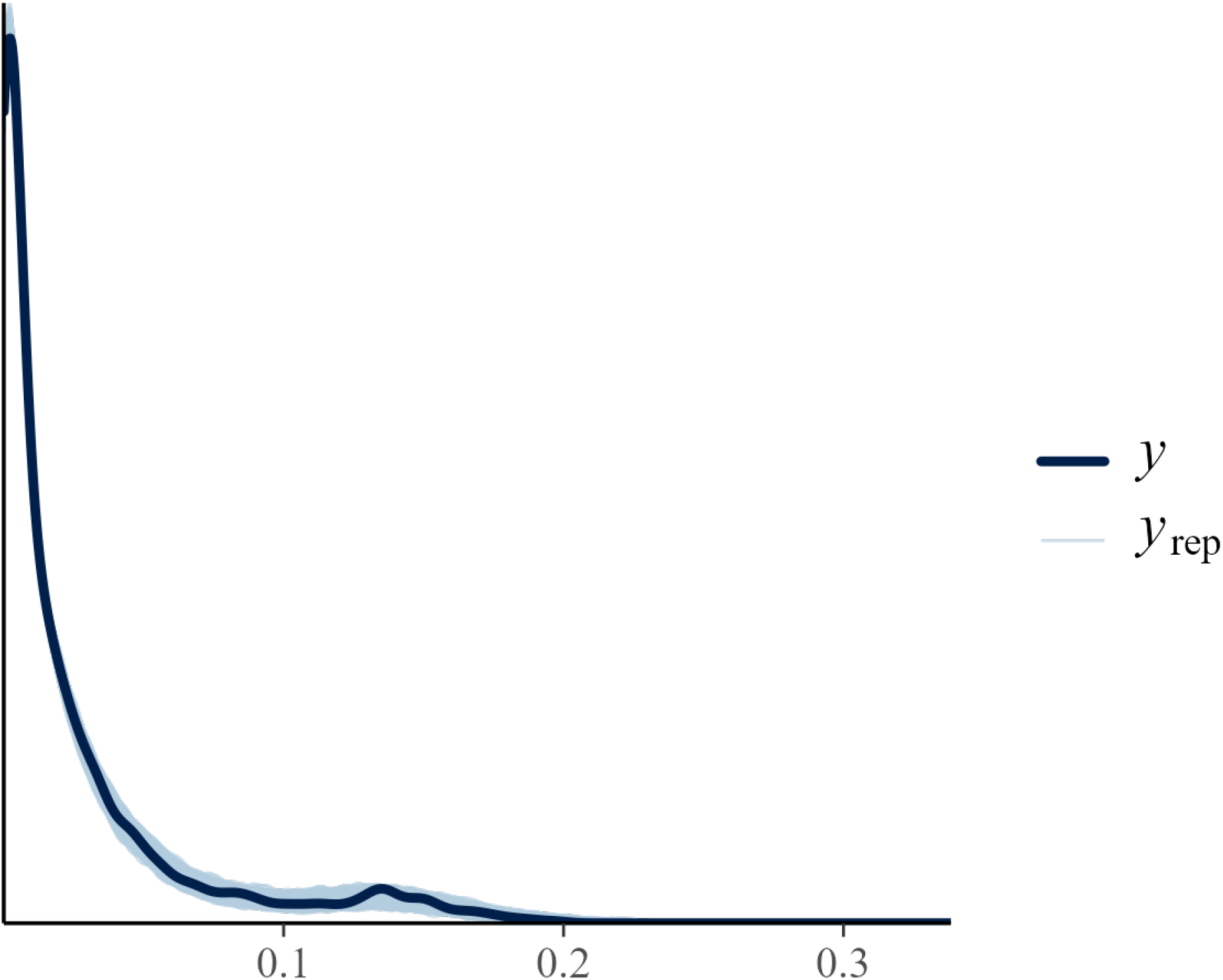
Density of the replicated probability of deaths overlaid by density of the observed probability of deaths shows that the model approximately reproduces the observed data. The dark blue line represents the density of the observed data *q*(*x, t, l*) labelled as *y* and the light blue lines represent the density of each replicate of the probability of death produced by the Bayesian model, labelled as *y*_*rep*_.

**Figure A2:**
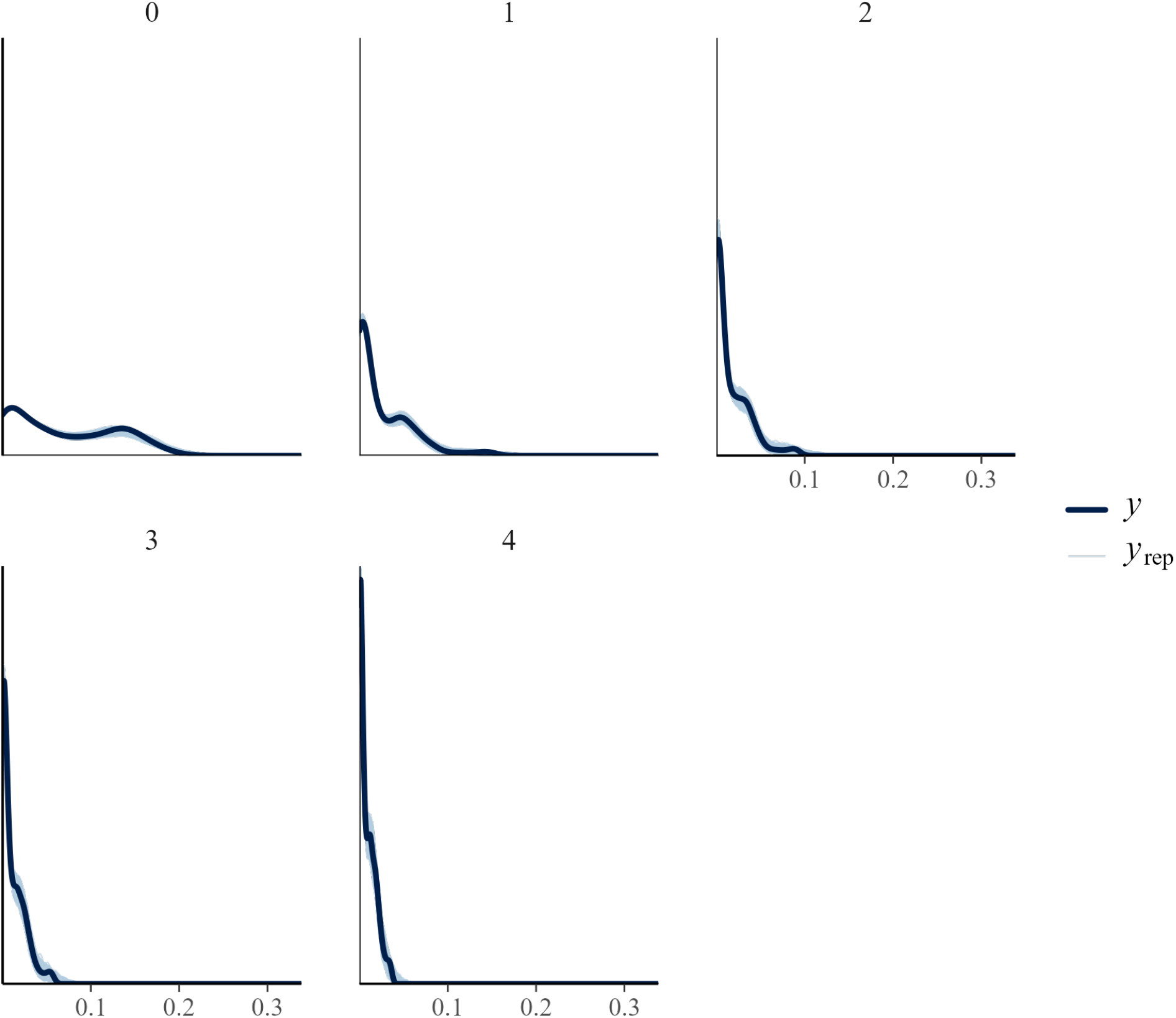
Density of the replicated probability of deaths overlaid by the density of observed probability of deaths by age indicates a good model fit. The dark blue line represents the density of the observed data *q*(*x, t, l*) labelled as *y* and the light blue lines represent the density of each replicate of the probability of death produced by the Bayesian model, labelled as *y*_*rep*_. The ages *x* ∈ [0, 4] are mentioned at the top of each panel.

**Figure A3:**
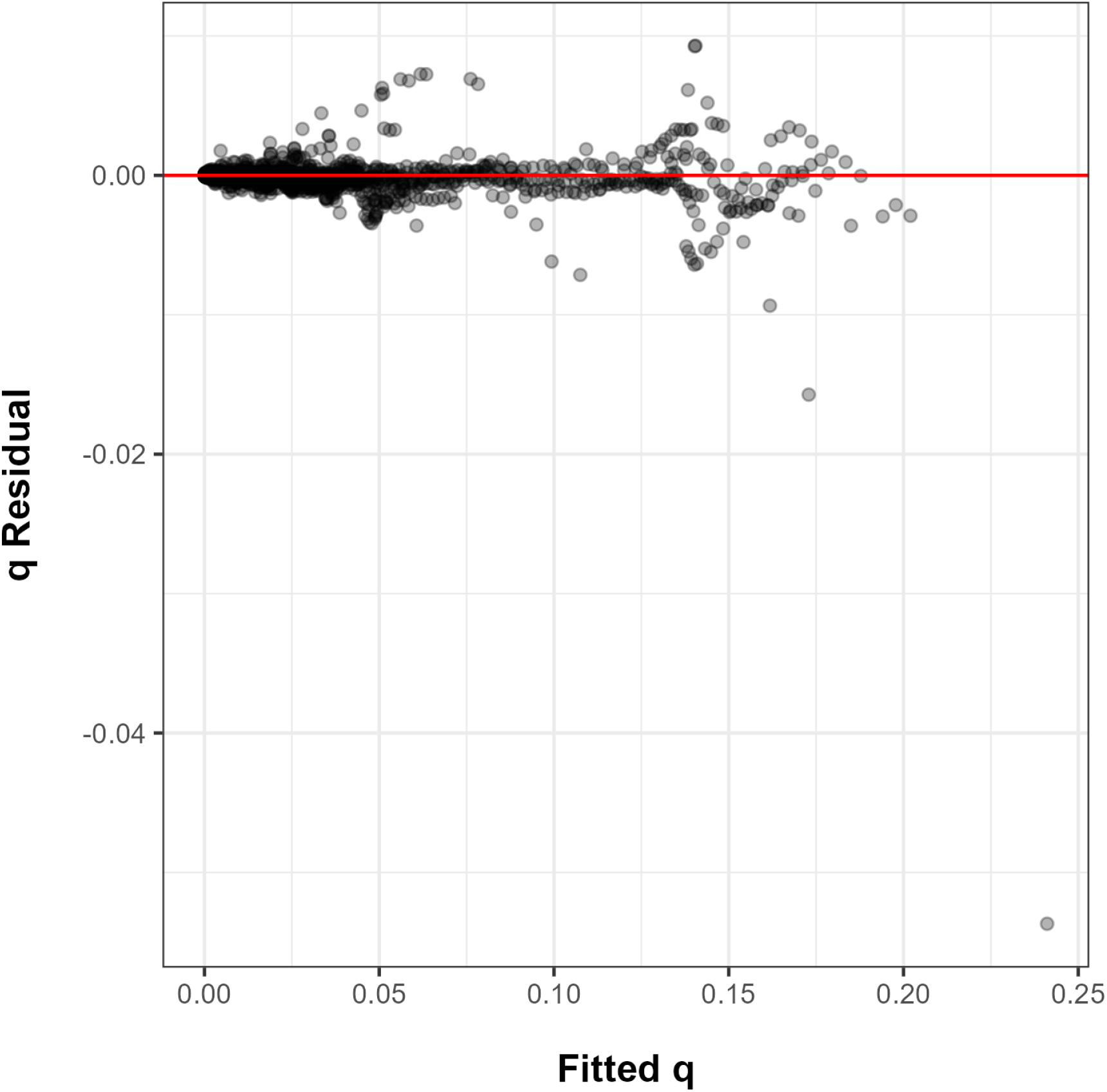
Residuals and the fitted probability of death shows a pattern indicating good fit of the model to the data, however, an outlier exists for one observation point. Almost all the residuals aligns closely around the horizontal line at zero and do not show unique pattern.

**Figure A4:**
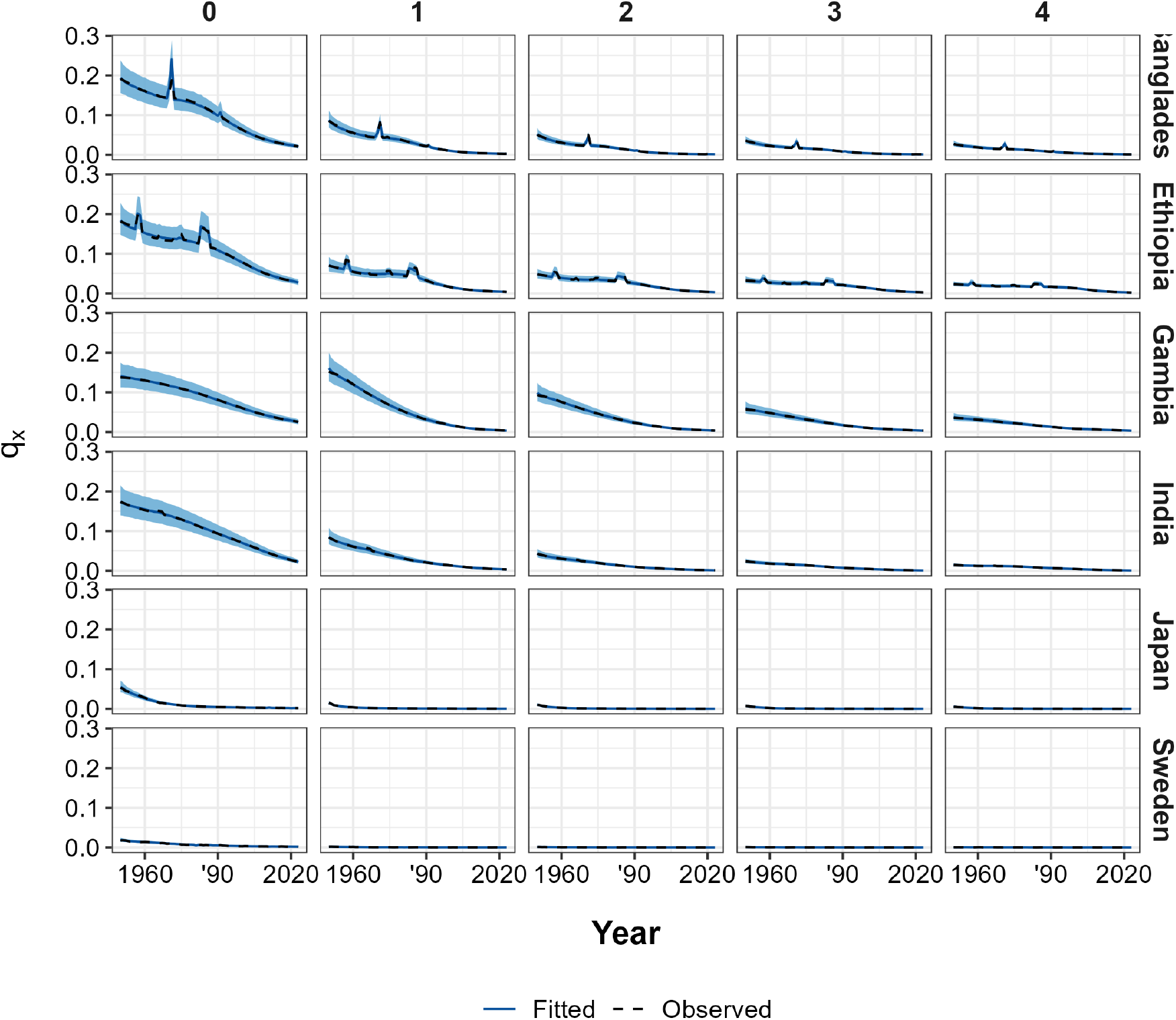
The median value of estimated probability of death presented with 95 percent uncertainty interval shows a good model fit to the observed data. The black dashed line indicates the observed probability of death and the thick blue line indicates the median of the posteriors of the estimated probability of death. The light blue band around the median estimates shows the 95 percent uncertainty interval.

**Figure A5:**
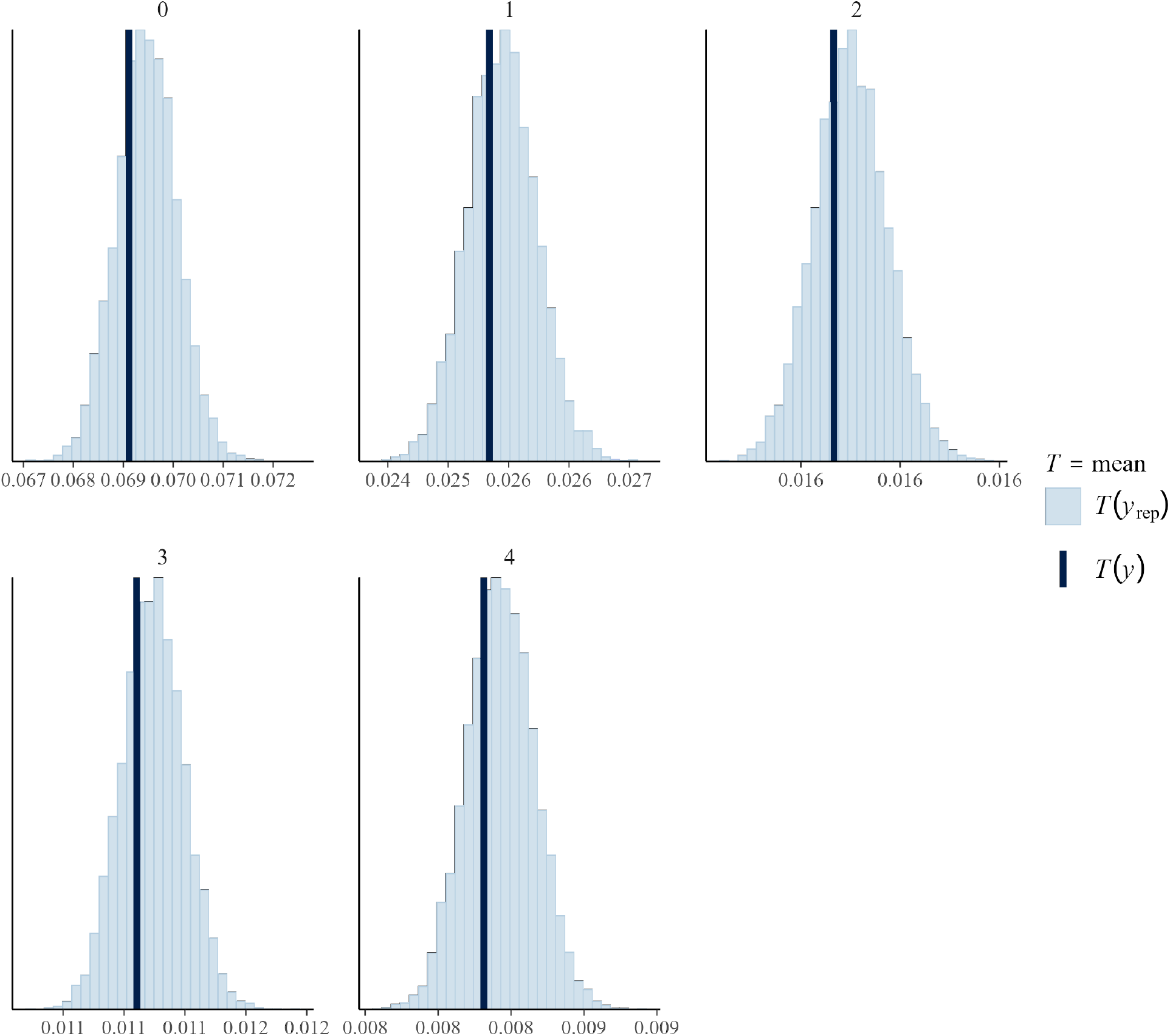
Mean of the replicated probability of deaths overlaid by the mean of the observed data by age indicates a good model fit. The histogram represents the mean of the probability of deaths for the replicates labelled as *y*_*rep*_. The dark blue line indicates the mean of the observed probability of death. The blue line at the centre of histograms for every age indicates a good model fit.

**Figure A6:**
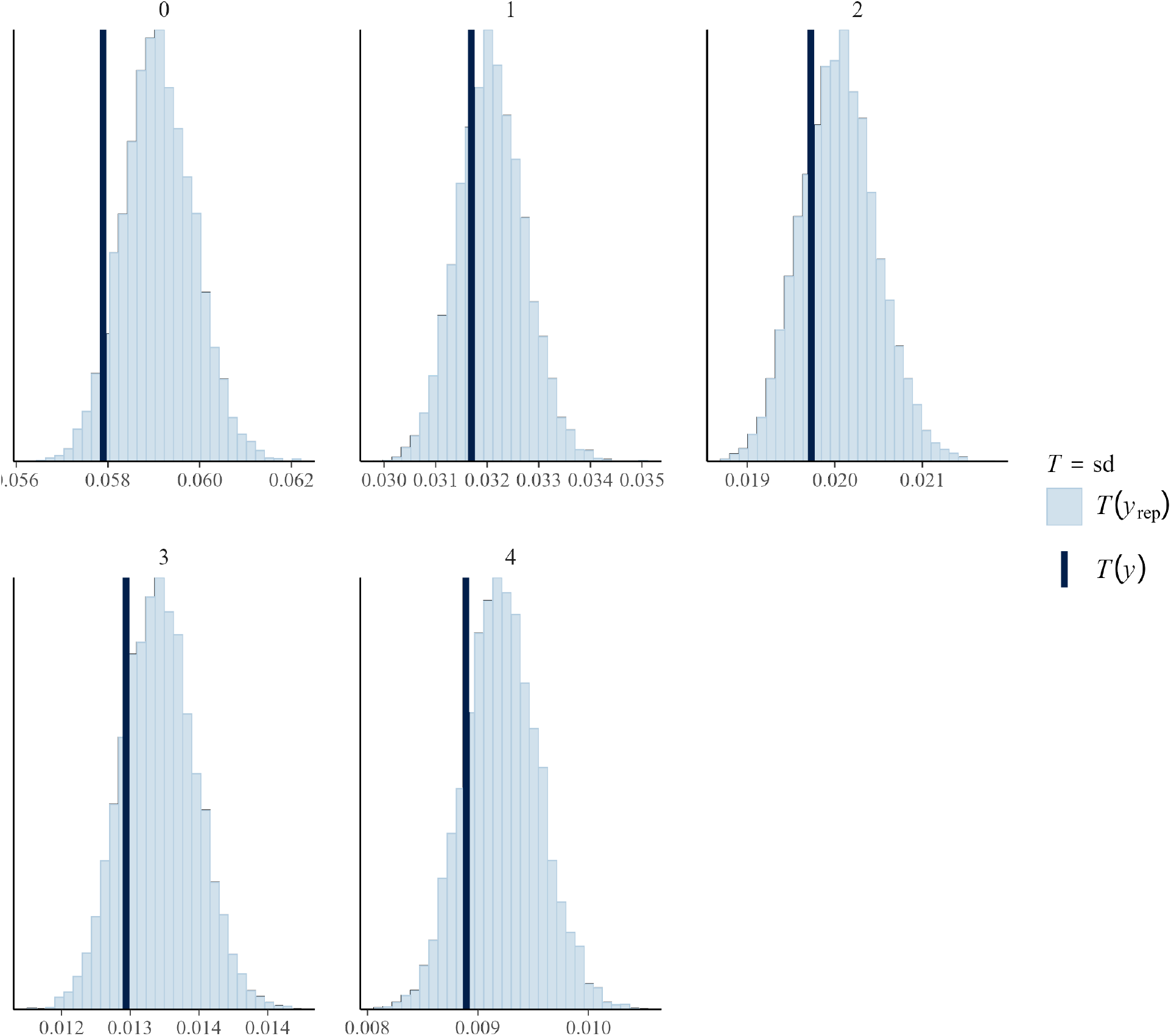
Standard deviations of the replicated probability of death overlaid by that of the observed data by age shows approximately a good model fit. The histograms shows the standard deviations for each replicates of the probability of death labelled as *y*_*rep*_ and the tick blue line represents the standard deviation for the observed data labelled as *y*. The standard deviations for all the ages but for age zero lies at the center of the histogram.

**Figure A7:**
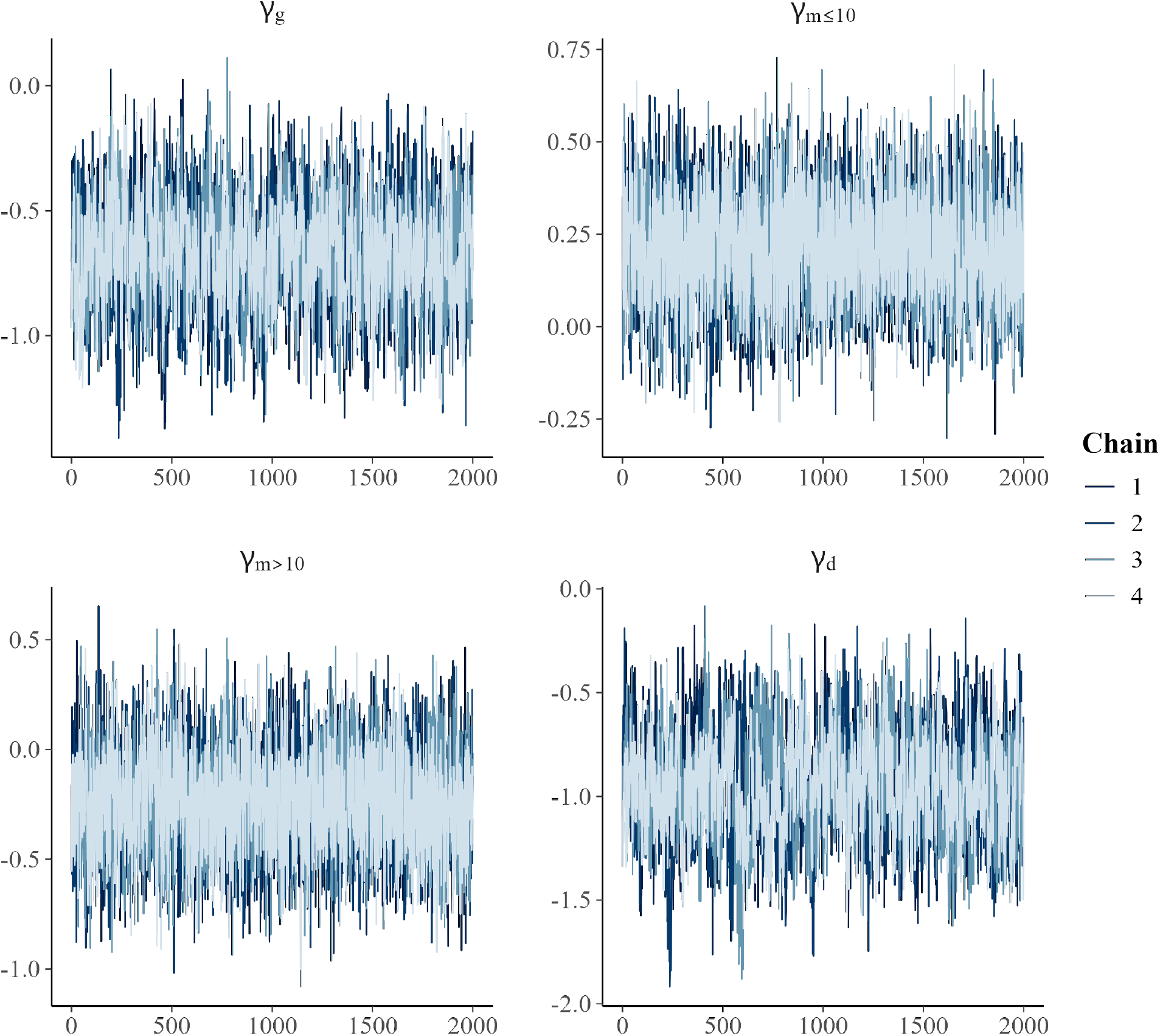
Trace plots for the slope coefficient of the probability of having a specific type of kin shows that the slope coefficients for the four chains converged. The samples for each chain presented in different shades of blue overlap considerably. The trace plots for the coefficients of grandmother *γ*_*g*_, mother *γ*_*d*_, older sister age more than 10 years *γ*_*m>*10_, and older sisters age less than or equal to 10 years *γ*_*m*≤10_ are lablled accordingly on top of each panel.

## Notes

### Competing Interest Statement

The authors have declared no competing interest.

